# Clathrin-associated carriers enable recycling through a kiss-and-run mechanism

**DOI:** 10.1101/2024.07.09.601372

**Authors:** Jiachao Xu, Yu Liang, Nan Li, Song Dang, Amin Jiang, Yiqun Liu, Yuting Guo, Xiaoyu Yang, Yi Yuan, Xinyi Zhang, Yaran Yang, Yongtao Du, Anbing Shi, Xiaoyun Liu, Dong Li, Kangmin He

## Abstract

Endocytosis and recycling control the uptake and retrieval of various materials, including membrane proteins and lipids, in all eukaryotic cells. These processes are crucial for cell growth, organization, function, and environmental communication. However, the mechanisms underlying efficient, fast endocytic recycling remain poorly understood. Here, by utilizing a biosensor and imaged-based screening, we uncover a novel recycling mechanism that couples endocytosis and fast recycling, which we name the clathrin-associated fast endosomal recycling pathway (CARP). Clathrin-associated tubulovesicular carriers containing clathrin, AP1, Arf1, Rab1, and Rab11, while lacking the multimeric retrieval complexes, are generated at subdomains of early endosomes, and then transported along actin to cell surfaces. Unexpectedly, the clathrin-associated recycling carriers undergo partial fusion with the plasma membrane. Subsequently, they are released from the membrane by dynamin and reenter cells. Multiple receptors utilize and modulate CARP for fast recycling following endocytosis. Thus, CARP represents a novel endocytic recycling mechanism with kiss-and-run membrane fusion.

## Main

Endocytosis is a fundamental and essential process of eukaryotic cells to internalize integral cell surface proteins as well as external materials such as nutrients and pathogens^1^. Once internalized, most materials are rapidly transported to early endosomes for endosomal sorting to the recycling or degradation pathways. Approximately 70–80% of internalized cell-surface proteins, including receptors, channels, transporters, and cell adhesion molecules, are recycled back to the plasma membrane^2, 3^. Endocytosis and recycling are essential for cell survival and homeostasis, and communication with the environment. The dysregulation of these processes is closely related to various human diseases, particularly cancer and neurodegenerative disorders^4, 5^. Although the dynamics and regulation of different endocytosis pathways have been well characterized, the pathways and associated mechanism(s) by which thousands of membrane proteins, including signaling receptors, are correctly and efficiently recycled back to the plasma membrane remain elusive.

Early endosomes are highly dynamic and pleomorphic structures with vacuoles (∼100-500 nm diameter) and multiple thin tubular extensions^6, 7^. As the first and primary sorting station of cells, early endosomes receive internalized cargoes and subsequently sort them either to the recycling pathways for transport back to the cell membrane, to late endosomes and lysosomes for degradation, or to the Golgi and the trans-Golgi network (TGN) for recycling and retrieval^2, 3, 8, 9^. Mammalian cells have developed several recycling routes including the relatively extensively studied “fast” and “slow” tubulovesicular recycling pathways. The “fast” recycling pathway, regulated mainly by Rab4 or Rab35, directly sorts internalized cargo from early endosomes back to the cell surface. In contrast, the “slow” recycling pathway, regulated mainly by Rab11, recycles cargo from the juxtanuclear early recycling compartments or recycling endosomes^6, 8^. Certain receptors, such as transferrin receptors (TfR), can utilize both the fast and slow recycling pathways for constitutive recycling while signaling receptors like the β2 adrenergic receptor (β2AR) of G protein-coupled receptors (GPCRs) require specific sequences (sorting signals) for stimulation-dependent fast recycling^10^. Nevertheless, the existence of endocytic recycling pathways beyond the canonical “fast” and “slow” recycling pathways remains largely unknown.

In our previous work, we introduced “coincidence detecting” sensors that selectively report the phosphoinositide composition of clathrin-associated structures^11^. In this study, we have further utilized the phosphatidylinositol-4,5-biphosphate (PI(4,5)P_2_) fluorescent sensor, which selectively binds to clathrin-coated structures physically connected to the plasma membrane. Through imaging based-screening, we have identified the clathrin-associated fast endosomal recycling pathway (CARP), which starts with the generation of clathrin-associated carriers from early endosomes, followed by partial fusion of these carriers with the plasma membrane, and subsequent scission and budding off from the plasma membrane.

## Results

### Detection of AP2-negative clathrin-associated carriers using the coincidence detecting PI(4,5)P_2_ sensor

Clathrin-mediated endocytosis begins with clathrin assembly on the plasma membrane to form clathrin-coated pits, a process initiated by clathrin and adaptor proteins, particularly the adaptor protein 2 (AP2) complex^12, 13^. PI(4,5)P_2_ is the most abundant phosphoinositide species located at the inner leaflet of the plasma membrane^14^. PI(4,5)P_2_ diffuses rapidly (approximately 0.3 μm^2^/s) and freely within the plasma membrane and connected membrane structures^15^. Using the clathrin-coated structure-specific PI(4,5)P_2_ sensor (EGFP-PH(PLCδ1)-Aux1), we successfully detected and tracked the existence and dynamics of PI(4,5)P_2_ associated with clathrin-coated pits and vesicles at the plasma membrane^11^. The essence of the coincidence detecting sensor lies in its simultaneous weak binding of clathrin by the clathrin-binding domain of auxilin1 (Kd of ∼100–200 μM) and PI(4,5)P_2_ by the PH domain of PLC81 (Kd of ∼2 μM)^16^ in membrane structures containing both clathrin and PI(4,5)P_2_. Neither the clathrin-binding domain nor the PI(4,5)P_2_-binding domain alone is adequate for effective recruitment to clathrin-coated structures^11^. The presence of both PI(4,5)P_2_ and clathrin in membrane structures is crucial as it provides binding sites for both domains, thereby achieving a sufficiently high affinity to recruit the coincidence detecting sensor^11^ (Fig. 1a).

**Fig. 1.**
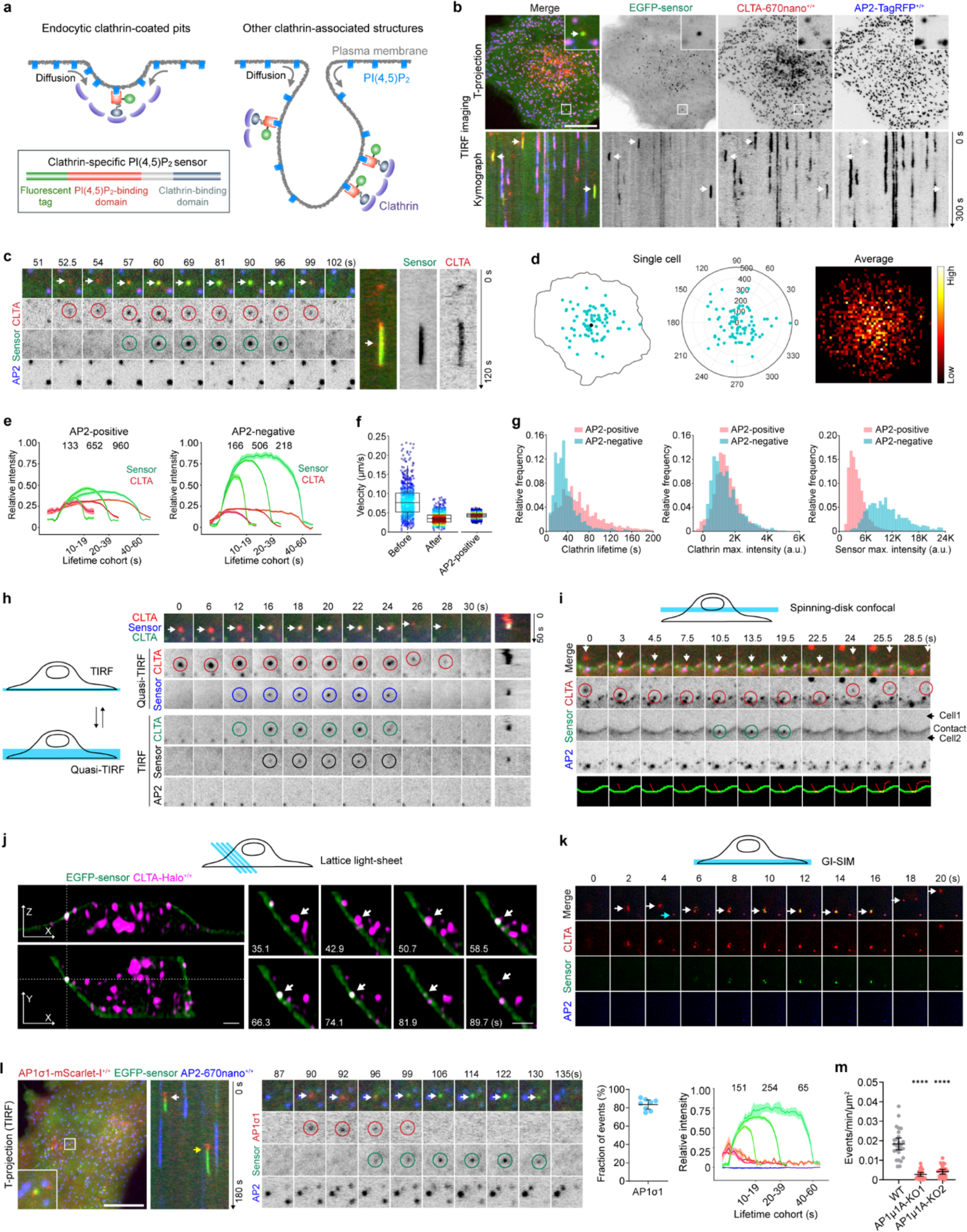
Identification of intracellular clathrin-associated carriers that transiently visit the plasma membrane. **a**, Lower left: Domain organization of the coincidence detecting PI(4,5)P_2_ sensor, consisting of the fluorescent tag, the PI(4,5)P_2_-binding domain, and the clathrin-binding domain of auxilin1. Top left: Diagram illustrating sensor association with endocytic clathrin-coated pits (adapted from^11^). Right: Diagram illustrating sensor association with intracellular clathrin-associated structures that are physically connected to the plasma membrane. The established membrane continuity enables the rapid diffusion of PI(4,5)P_2_ from the plasma membrane to the clathrin-associated structures. **b**, Bottom surfaces of gene-edited AP2-TagRFP^+/+^ CLTA-670nano^+/+^ cells stably expressing the coincidence detecting PI(4,5)P_2_ sensor EGFP-PH(PLC81)-Aux1 (EGFP-sensor). Cells were imaged every 1.5 s by TIRF microscopy. The maximum-intensity time projection (T-projection) and kymographs of a representative time series are shown. The AP2-negative clathrin-positive events that recruited the EGFP-sensor (AP2-negative events) are highlighted by arrows. SUM159 cells were used unless otherwise noted. **c**, Representative montage (left) and kymographs (right) showing the dynamics of an AP2-negative event at the plasma membrane. **d**, Left: AP2-negative events in a time series were identified and displayed (dots) within the cell boundary. Middle: AP2-negative events in the circularly normalized cell. Right: Spatial frequency distribution of events from 21 cells. **e**, Averaged fluorescence intensity traces of AP2-positive endocytic events (left) and AP2-negative events (right) from 21 cells (mean ± SEM), grouped by cohorts according to the lifetimes of clathrin (intensity cohorts). Numbers of analyzed traces are shown above each cohort. **f**, Distributions of the velocity of AP2-negative events before and after recruiting the EGFP-sensor, and the AP2-positive endocytic events (median with interquartile range, n = 21 cells). **g**, Left and middle: Distributions of lifetimes and maximum (max.) fluorescence intensities of clathrin from AP2-positive endocytic events and AP2-negative events. Right: Distributions of max. fluorescence intensities of EGFP-sensor from the events (10969 and 1209 events from 21 cells). **h**, CLTA-Halo_+/+_ AP2-TagRFP^+/+^ cells stably expressing EGFP-sensor were labeled with the JFX_650_-HaloTag ligand, and then imaged every 2 s by alternating the incident angles between the critical angle (TIRF) and the angle smaller than the critical angle (Quasi-TIRF). The montages and kymographs show the detection of the AP2-negative clathrin-positive carrier by Quasi-TIRF first and then TIRF, followed by the recruitment of the sensor at the plasma membrane. The schematic representation of sequential TIRF and Quasi-TIRF illumination is depicted on the left. **i**, CLTA-Halo^+/+^ AP2-TagRFP^+/+^ cells stably expressing EGFP-sensor were labeled with the JFX_650_-HaloTag ligand, and then imaged at the middle plane every 1.5 s by spinning-disk confocal microscopy. The montage shows the dynamic movement of an intracellular clathrin-positive carrier to the plasma membrane, followed by the recruitment of the sensor and the subsequent departure of the carrier from the plasma membrane. The tracks of the clathrin-positive carrier (red) and the segmented plasma membrane between the two cells (green) are displayed at the bottom. **j**, CLTA-Halo^+/+^ AP2-TagRFP^+/+^ cells stably expressing EGFP-sensor were labeled with the JFX_650_-HaloTag ligand and then imaged in 3D every 3.9 s by lattice light-sheet microscopy. Left: The XZ and XY optical sections of the cell at a single time point (66.3 s), capturing a fusion event. Right: Montage showing the dynamic movement of an intracellular clathrin-positive carrier to the plasma membrane, followed by the sudden recruitment of the sensor at the plasma membrane (arrows). **k**, CLTA-TagRFP^+/+^ AP2-Halo^+/+^ cells stably expressing EGFP-sensor were labeled with the JF_646_-HaloTag ligand, and then imaged every 2 s by GI-SIM near the bottom surface. The representative montage shows the dynamic movement of an AP2-negative clathrin-positive carrier before, during, and after recruiting the sensor (white arrows). The blue arrow highlights an endocytic event that is triple-positive for AP2, clathrin, and sensor. **l**, AP1σ1-mScarlet-I^+/+^ AP2-670nano^+/+^ cells stably expressing EGFP-sensor were imaged by every 1 s TIRF microscopy. Left: T-projection and kymograph of a time series. Middle: Montage of the event indicated by the yellow arrow in the kymograph. Right: The fraction of AP2-negative sensor-positive events that showed AP1 recruitment (mean ± 95% confidence interval (CI)), as well as the intensity cohorts of AP1 (red), sensor (green), and AP2 (blue) (mean ± SEM; n = 8 cells). **m**, AP2-TagRFP^+/+^ CLTA-670nano^+/+^ cells stably expressing EGFP-sensor underwent knockout of AP1μ1A expression using two different sgRNAs, and then imaged every 1 s by TIRF microscopy. The plots show the frequency of the AP2-negative clathrin-positive events that recruited the EGFP-sensor from 26 wild-type, 24 AP1μ1A-KO1, and 24 AP1μ1A-KO2 cells (mean ± 95% CI). |Statistical analysis was performed using the ordinary one-way ANOVA with Tukey’s multiple comparisons test in **m**; \*\*\*\**P* < 0.0001. Scale bars, 10 μm in **b** and **l**, and 2 μm in **j**.

We utilized the EGFP-PH(PLC81)-Aux1 sensor for triple-color live-cell imaging using spinning-disk confocal microscopy of SUM159 cells, which were dually genome-edited to express AP2-TagRFP^+/+^ (TagRFP fused to the C-terminus of the σ2 subunit of AP2) and CLTA-Halo^+/+^ (HaloTag fused to the C-terminus of clathrin light chain A and stained with the JFX_650_-HaloTag ligand before imaging) (Extended Data Fig. 1a). Consistent with the enrichment of PI(4,5)P_2_ at the plasma membrane, the coincidence detecting PI(4,5)P_2_ sensor EGFP-PH(PLCδ1)-Aux1 was recruited specifically to clathrin-associated structures at the plasma membrane and did not associate with intracellular clathrin-coated structures in endosomes or the trans-Golgi network (TGN)^11^ (Extended Data Fig. 1c). Unexpectedly, in addition to the anticipated endocytic coated pits triple-positive for clathrin, AP2, and the sensor, we noticed a population of sensor-positive but AP2-negative clathrin-associated structures located at the plasma membrane (Extended Data Fig. 1c).

To characterize the dynamics of the unexpected AP2-negative structures found at the plasma membrane, we stably expressed the EGFP-PH(PLCδ1)-Aux1 sensor at a low level in the dually genome-edited SUM159 cells expressing AP2-TagRFP^+/+^ and CLTA-670nano^+/+^ (miRFP670nano) (Extended Data Fig. 1b) and imaged the cells at the basal plasma membrane by triple-color total internal reflection fluorescence (TIRF) microscopy. Time-lapse TIRF imaging also unveiled the presence of a population of short-lived clathrin-associated carriers that were positive for the EGFP-PH(PLCδ1)-Aux1 sensor but negative for AP2 (Fig. 1b,c and Supplementary Video 1). The instantaneous appearance of short-lived clathrin-labeled structures in the TIRF imaging field has also been noted in previous studies^17^. Notably, the clathrin-positive AP2-negative carriers displayed fast movement beneath the plasma membrane, temporarily stalled, and then quickly recruited a significant amount of the sensor. After staying approximately 5-50 s at the plasma membrane with the sensor, the clathrin-positive AP2-negative carriers moved away from the plasma membrane without the sensor (Fig. 1b,c and Supplementary Video 1). These events occurred primarily near the central region of the plasma membrane (Fig. 1d). In contrast to AP2-positive endocytic clathrin-coated pits, which exhibited a gradual increase in the fluorescence signal of clathrin reflecting clathrin coat assembly, the clathrin-positive AP2-negative carriers tended to appear promptly (Fig. 1e,f) and exhibited a shorter residence time at the plasma membrane (Fig. 1g). They also showed a higher fluorescence signal of the EGFP-PH(PLCδ1)-Aux1 sensor compared to the endocytic carriers (Fig. 1e,g). Knockout of the AP2 σ2 subunit in the genome-edited cells expressing AP2-TagRFP^+/+^ CLTA-Halo^+/+^ inhibited transferrin endocytosis (Extended Data Fig. 1d) but did not affect the sudden appearance of the clathrin-associated carriers that recruited the sensor at the plasma membrane (Extended Data Fig. 1e,f). Thus, these AP2-negative clathrin-positive carriers that recruited the EGFP-PH(PLCδ1)-Aux1 sensor are distinct from and less likely to be endocytic carriers.

To demonstrate that the recruitment of the coincidence detecting sensor EGFP-PH(PLCδ1)-Aux1 to clathrin-associated structures relies on its binding to PI(4,5)P_2_ at the plasma membrane, we acutely depleted PI(4,5)P_2_ by rapid recruitment of the inositol 5-phosphatase module of OCRL (5-ptaseOCRL) from the cytosol to the plasma membrane. PI(4,5)P_2_ depletion from the plasma membrane readily prevented the recruitment of the sensor to both AP2-positive endocytic structures and AP2-negative clathrin-associated structures (Extended Data Fig. 1g). In addition to employing cells genome-edited to fuse a fluorescent tag to the C-terminus of the σ2 subunit of AP2, we established stable cell lines expressing the internally TagRFP-tagged α subunit of AP2, as previously described^18^. The EGFP-PH(PLCδ1)-Aux1 sensor is strongly recruited to these AP2-negative motile clathrin-associated carriers after they stall at the plasma membrane of cells stably expressing α-TagRFP-AP2 (Extended Data Fig. 1h). Moreover, by replacing PH(PLCδ1) with another PI(4,5)P_2_-binding domain Tubby_c_ (C-terminal domain from Tubby; Kd >2 μM)^16^, the new coincidence detecting sensor EGFP-Tubby_c_-Aux1 was similarly recruited to AP2-positive endocytic structures and the AP2-negative clathrin-associated carriers (Extended Data Fig. 1i and Supplementary Video 2). Introducing point mutations into the lipid-binding domain of PH(PLCδ1) or Tubby_c_ to abolish PI(4,5)P_2_ binding prevented the recruitment of the sensors to clathrin-associated structures (Extended Data Fig. 1j,k). In addition to the coincidence detecting sensor, we fused PH(PLCδ1) or Tubby_c_ with the bright fluorescent protein mNeonGreen and observed the anticipated uniform distribution across the plasma membrane. Occasionally, we observed the weak recruitment of mNeonGreen-PH(PLCδ1) or mNeonGreen-Tubby_c_ to certain brighter AP2-negative clathrin-associated structures when they briefly paused around the plasma membrane (Extended Data Fig. 1l). Furthermore, the AP2-negative clathrin-associated carriers that transiently recruited the EGFP-PH(PLCδ1)-Aux1 sensor were also observed in two additional cell lines (human U2OS cells and green monkey COS7 cells) (Extended Data Fig. 1m,n).

### The AP2-negative clathrin-associated structures are intracellular carriers that transiently visit the plasma membrane

Since the AP2-negative clathrin-associated carriers exhibit fast movement before stalling and recruiting the coincidence detecting sensors at the plasma membrane, we then investigated whether these structures are intracellular carriers that transiently visit the plasma membrane. To characterize the three-dimensional (3D) dynamics of the AP2-negative clathrin-associated carriers, we employed variable-angle TIRF microscopy, spinning-disk confocal microscopy, and lattice light-sheet microscopy. Alternating the incident angles of TIRF between the critical angle (to limit imaging depth at the plasma membrane) and the angle smaller than the critical angle (to increase imaging depth into the cytoplasm, Quasi-TIRF) enables the exploration of the Z-dimensional movement of intracellular carriers docking at the plasma membrane^19^. The AP2-negative clathrin-associated carriers observed in TIRF were consistently first detected in Quasi-TIRF (Fig. 1h), confirming that the carriers are not assembled at the plasma membrane but instead are preformed intracellular structures that migrate to the plasma membrane. The sensor was recruited only when the intracellular carriers approached the plasma membrane. Subsequently, the clathrin-associated carriers departed and moved away from the plasma membrane (Fig. 1h). By imaging the cells at the middle plane by spinning-disk confocal microscopy and in 3D by lattice light-sheet microscopy, the intracellular movement and subsequent docking of clathrin-associated carriers at the plasma membrane, followed by sensor recruitment, can be directly observed (Fig. 1i,j and Supplementary Video 3). Live-cell grazing incidence structured illumination microscopy (GI-SIM) also enables the visualization of the rapid movement and subsequent docking of these nonspherical clathrin-associated structures around the plasma membrane (Fig. 1k and Supplementary Video 4).

The next question is what are these intracellular carriers that contain clathrin? Adaptor protein AP1 is involved in the formation of intracellular clathrin-coated structures originating from endosomes and TGN^20^. By imaging the EGFP-PH(PLCδ1)-Aux1 sensor in SUM159 cells genome-edited to express AP1-mScarlet-I^+/+^ (mScarlet-I fused to the C-terminus of the σ1 subunit of AP1) (Extended Data Fig. 2a), we readily observed the membrane docking of intracellular AP2-negative AP1-positive carriers, followed by the disappearance of AP1 signal and rapid recruitment of the sensor (Fig. 1l and Supplementary Video 5). The membrane docking and sensor recruitment were similarly observed with the AP1-positive carriers labeled by the μ1A subunit of AP1 (Extended Data Fig. 2b). Remarkably, AP1 is associated with nearly all the AP2-negative carriers before sensor recruitment (Fig. 1l and Extended Data Fig. 2b). Knockout of AP1 significantly inhibited the appearance of the clathrin-positive carriers that recruited the sensor at the plasma membrane (Fig. 1m and Extended Data Fig. 2c), indicating the essential role of AP1 in clathrin-associated carrier generation.

### Intracellular clathrin-associated carriers undergo fusion with the plasma membrane

While the effective recruitment of the coincidence detecting PI(4,5)P_2_ sensor depends on its binding to both clathrin and the plasma membrane-localized PI(4,5)P_2_, the intracellular clathrin/AP1-associated carriers contain clathrin but not PI(4,5)P_211_ (Extended Data Fig. 1c). Thus, the recruitment of the sensor to the clathrin/AP1-associated carriers during their period of stalling at the plasma membrane suggests that the carriers can transiently fuse with the plasma membrane. The continuity of the membrane established during membrane fusion would allow the carriers to rapidly acquire PI(4,5)P_2_ through the diffusion of PI(4,5)P_2_ from the plasma membrane, thereby inducing the rapid recruitment of the coincidence detecting sensor (Fig. 1a). The docking and membrane fusion of exocytic carriers with the plasma membrane is mediated by the exocyst complex and soluble *N*-ethylmaleimide-sensitive factor attachment protein receptor (SNARE) proteins^21^ (Fig. 2a). To demonstrate that the abrupt recruitment of the sensor by clathrin-associated carriers reflects membrane fusion, we monitored the dynamic association as well as functions of the exocyst and SNARE proteins during the fusion of clathrin-associated carriers with the plasma membrane. The exocyst, an evolutionarily conserved octameric complex, plays a crucial role in tethering exocytic vesicles to the plasma membrane before SNARE-mediated fusion^22, 23^. The exocyst complex consists of Sec3, Sec5, Sec6, Sec8, Sec10, Sec15, Exo70 and Exo84 subunits^23^. By generating genome-edited cell lines with endogenous Exo70, Sec3, and Sec5 tagged with fluorescent proteins (Extended Data Fig. 2d-f), we observed that endogenous Exo70, Sec3, and Sec5 were invariably recruited before the abrupt appearance of the sensor signal at the plasma membrane (Fig. 2b,c, Extended Data Fig. 2g, Supplementary Videos 6 and 7). Notably, almost all the AP2-negative carriers that transiently recruited the sensor exhibited recruitment of endogenous Sec3, Exo70, and Sec5 (Fig. 2d). Other ectopically expressed exocyst components were also robustly recruited immediately before sensor recruitment (Extended Data Fig. 2h,i).

**Fig. 2.**
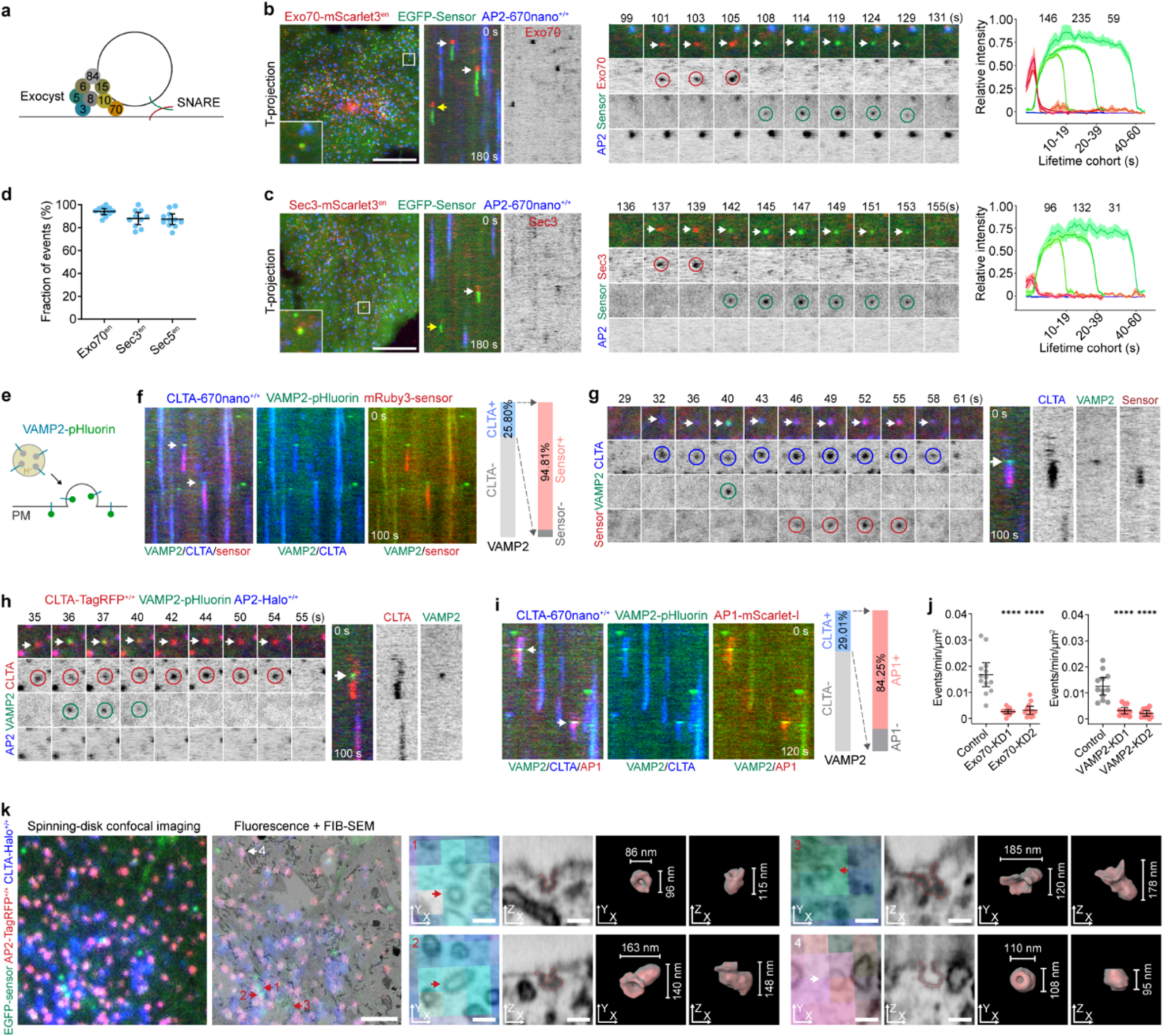
Fusion of the intracellular clathrin-associated carriers with the plasma membrane. **a**, Diagram of the assembly of exocyst and SNARE complexes during the tethering and fusion of exocytic vesicles. **b**, Genome-edited AP2-670nano^+/+^ and Exo70-mScarlet3 (pool) cells stably expressing EGFP-sensor were imaged every 1 s by TIRF microscopy. Left: T-projection and kymographs of a time series. Middle: Montage of the event indicated by the yellow arrow in the kymograph. Right: The intensity cohorts of Exo70 (red), sensor (green), and AP2 (blue) of the AP2-negative sensor-positive events (mean ± SEM; n = 12 cells). **c**, Genome-edited AP2-670nano^+/+^ and Sec3-mScarlet3 (pool) cells stably expressing EGFP-sensor were imaged and displayed as in (**b**). Intensity cohorts from 14 cells. **d**, The fraction of AP2-negative sensor-positive events that showed endogenous Exo70, Sec3, or Sec5 recruitment (mean ± 95% CI; n = 11, 10, and 11 cells). **e**, Diagram illustrating the dramatic increase in the fluorescence signal of pHluorin (attached to the C-terminal luminal part of VAMP2) during the fusion of an exocytic vesicle with the plasma membrane. **f**, CLTA-670nano^+/+^ cells stably expressing VAMP2-pHluorin were transiently transfected with mRuby3-sensor, and then imaged every 1 s by TIRF microscopy. Kymographs show the sudden recruitment of mRuby3-sensor to the VAMP2/clathrin-positive fusion events (arrows). The fraction of VAMP2-positive exocytosis events that contained clathrin and the fraction of VAMP2/clathrin-positive events that recruited the sensor are shown on the right (1868 events from 8 cells). **g**, Representative montage and kymographs showing the recruitment of mRuby3-sensor to a clathrin- associated carrier after the transient burst of VAMP2-pHluorin at the plasma membrane. **h**, CLTA-TagRFP^+/+^ AP2-Halo^+/+^ cells stably expressing VAMP2-pHluorin were labeled with the JFX_650_- HaloTag ligand and then imaged every 1 s by TIRF microscopy. The representative montage and kymographs show the burst recruitment of VAMP2 to an AP2-negative clathrin-positive fusion event at the plasma membrane. **i**, CLTA-670nano^+/+^ cells stably expressing VAMP2-pHluorin were transiently transfected with AP1σ1- mScarlet-I, and then imaged every 1 s by TIRF microscopy. Kymographs show the recruitment of AP1 to the VAMP2/clathrin-positive events (arrows). The fraction of VAMP2-positive exocytosis events that contained clathrin and the fraction of VAMP2/clathrin-positive events that contained AP1 are shown on the right (2627 events from 11 cells). **j**, AP2-TagRFP^+/+^ CLTA-670nano^+/+^ cells stably expressing EGFP-sensor were treated with control siRNA or two different siRNAs targeting either Exo70 (left) or VAMP2 (right), and then imaged every 1 s by TIRF microscopy. The plots show the frequency of the AP2-negative clathrin-positive events that recruited the EGFP-sensor from 13, 13, and 13 cells (left) and 12, 12, and 12 cells (right) (mean ± 95% CI). **k**, Correlative spinning-disk confocal fluorescence imaging with FIB-SEM of AP2-TagRFP^+/+^ CLTA- Halo^+/+^ cells stably expressing EGFP-sensor. The cells were stained with the JF_646_-HaloTag ligand, imaged by spinning-disk confocal microscopy, and then imaged volumetrically by FIB-SEM. The left two panels show the fluorescence image near the cell bottom surface and the overlay of the fluorescence and FIB-SEM images. Arrows highlight clathrin and sensor-positive but AP2-negative structures (1-3) and clathrin, sensor, and AP2 triple-positive endocytic structures (4). The right panels present enlarged views of the numbered structures, segmentation of the numbered structures from the connected plasma membrane, and 3D reconstructions of the segmented structures. Statistical analysis was performed using the ordinary one-way ANOVA with Tukey’s multiple comparisons test; \*\*\*\**P* < 0.0001. Scale bars, 10 μm in **b** and **c**; 2 μm (left panel) and 0.1 μm (right panels) in **k**.

Vesicle-associated membrane protein 2 (VAMP2), a core component of the SNARE complex, resides on the membrane of intracellular trafficking vesicles and facilitates their fusion with the plasma membrane^24^. The pH-sensitive fluorescent protein pHluorin attached to VAMP2 is a widely used tool for identifying single vesicle exocytosis events using TIRF microscopy^22, 25^. pHluorin is quenched in the acidic environment of exocytic vesicles but shows a dramatic increase in fluorescence when exposed to the neutral extracellular environment during exocytic vesicle fusion with the plasma membrane^26^ (Fig. 2e). The typical vesicle fusion events observed under TIRF imaging show a sudden burst in the fluorescence of VAMP2- pHluorin at the plasma membrane, followed by its lateral dispersal to the surrounding plasma membrane^22, 25, 26^. By imaging the PH(PLCδ1)-Aux1 sensor along with VAMP2-pHluorin, we observed the successive appearance of clathrin signal, a burst signal of VAMP2-pHluorin, and then the emergence of the sensor signal at the plasma membrane (Fig. 2f,g). We found that a fraction of VAMP2-marked exocytosis events contained clathrin, while almost all clathrin-positive exocytosis events recruited the EGFP-PH(PLCδ1)- Aux1 sensor (Fig. 2f). The AP2-negative clathrin-positive exocytosis events were similarly observed without the expression of the PH(PLCδ1)-Aux1 sensor (Fig. 2h and Supplementary Video 8). Furthermore, consistent with the results obtained with the sensor, by imaging AP1-mScarlet-I together with VAMP2- pHluorin in the CLTA-670nano^+/+^ cells, we found that over 80% of the clathrin-positive exocytosis events also contained AP1 (Fig. 2i).

To further confirm that the recruitment of the coincidence detecting sensor relies on the fusion of clathrin-associated carriers with the plasma membrane, we knocked down the expression of Exo70 or VAMP2. Notably, interfering with vesicle fusion with the plasma membrane by depleting either Exo70 or VAMP2 markedly inhibited the fusion events described above (Fig. 2j and Extended Data Fig. 2j,k). Taken together, these results demonstrate that intracellular clathrin-containing structures can directly fuse with the plasma membrane, and the sudden appearance of the AP2-negative PH(PLCδ1)-Aux1 sensor-positive signal can serve as a simple yet reliable marker for identifying the fusion of intracellular clathrin-associated carriers with the plasma membrane (fusion events).

To capture the ultrastructure of clathrin-associated carriers during membrane fusion, we first used spinning-disk confocal microscopy to identify clathrin-positive structures at the plasma membrane that were positive for the sensor but negative for AP2. Subsequently, we utilized focused ion beam scanning electron microscopy (FIB-SEM) to reveal the ultrastructure of the identified structures (Extended Data Fig. 2l,m). In comparison to the smaller and more spherical-shaped AP2-positive endocytic structures, the relatively large and irregular AP2-negative structures tended to exhibit elliptical or tubular shapes (Fig. 2k). Interestingly, over 90% of the AP2-negative structures identified in fluorescence imaging (55 out of 60 events from 5 cells) were observed to be connected to the plasma membrane through a distinct opening pore facing the outside of the cell under FIB-SEM imaging (Fig. 2k), further confirming the fusion of the structure with the plasma membrane.

### Clathrin-associated recycling carriers are distinct from other reported clathrin-associated structures

The intracellular clathrin-associated carriers just described are quite different from other previously reported AP2-negative clathrin-associated structures, including “gyrating clathrin” structures^27, 28^, clathrin- associated post-Golgi vesicles^29^, and AP2-independent clathrin-coated endocytic structures^30^. “Gyrating clathrin” structures are characterized by their localized highly dynamic gyrating movement (≥ 3 μm/s) beneath the plasma membrane and are labeled by the Arf6(Q67L) but not AP1^27, 28^. The clathrin-associated carriers we identified did not exhibit the gyrating behavior and contained AP1 but not Arf6(Q67L) (Extended Data Fig. 3a). Moreover, overexpression of Arf6(Q67L) strongly stimulated the generation of “gyrating clathrin” structures^28^ but did not affect the generation of the clathrin-associated carriers we identified (Extended Data Fig. 3b). Previous studies have also indicated the involvement of clathrin in the secretion of post-Golgi vesicles^29^. However, by tracking the trafficking of post-Golgi vesicles using the thermosensitive VSVG (VSVGts)^31^, which is retained in the ER at 40 ℃ and redistributed to the Golgi and then cell surface upon a temperature shift to 32 ℃ (Extended Data Fig. 3c), we did not observe a close association between the PH(PLCδ1)-Aux1 sensor-recruited fusion events and these newly released VSVGts (15-20 min) at the plasma membrane (Extended Data Fig. 3d). By monitoring the trafficking of TfR1-containing post-Golgi vesicles with the “retention using selective hooks” (RUSH) system^32^, which relies on the synchronous release of the streptavidin-binding peptide (SBP)-fused TfR1 from the ER-localized streptavidin-fused hook protein upon the addition of biotin (Extended Data Fig. 3e), we further verified that the sensor-recruited fusion events are not closely related to the newly released TfR-containing post-Golgi vesicles (15-20 min) at the plasma membrane (Extended Data Fig. 3f). However, as released VSVGts and TfR1 reached trafficking equilibrium between the plasma membrane and endosomes (∼30 min), they began to exhibit a stronger association with the intracellular clathrin-associated carriers (Extended Data Fig. 3d,f). Furthermore, the fusion events we identified did not contain neuropeptide Y (NPY), a marker for secretory vesicles (Extended Data Fig. 3g). Previous studies have also suggested the existence of the AP2-independent clathrin-coated endocytic structures, which contains and relies on three endocytic adaptor proteins: Eps15, Eps15L1 (Eps15R), and Epsin1^30^. By generating genome-edited cell lines with endogenous Eps15, Eps15R, and Epsin1 tagged with fluorescent proteins, we found that the AP2- negative clathrin-associated carriers that recruited the PH(PLCδ1)-Aux1 sensor did not contain endogenous Eps15, Eps15R, or Epsin1 (Extended Data Fig. 3h-j).

Therefore, the above results demonstrate that the intracellular clathrin-associated carriers we identified are distinct from G-clathrin structures, post-Golgi secretory vesicles, and AP2-independent clathrin-coated endocytic structures. Given that the carriers we identified can mediate the recycling of endocytosed receptors back to the plasma membrane (shown in Fig. 8 and Extended Data Fig. 10), the term “clathrin-associated recycling carrier” is used in the following text to distinguish them from other clathrin-associated structures. The intracellular clathrin-associated recycling carriers that transiently fuse with the plasma membrane may represent a previously uncharacterized membrane trafficking pathway toward the plasma membrane.

### Imaging-based screening of biomolecules associated with CARP

The intricate intracellular trafficking process is regulated by a variety of trafficking machinery proteins, including Rab and Arf GTPases, coat proteins and their adaptors, cytoskeleton and motor proteins, tethering factors, and SNARE proteins^6, 8^. To characterize the molecular components and properties of the clathrin-associated recycling carriers, we conducted TIRF microscopy-based imaging screening for proteins that potentially associate with the carriers at different stages of trafficking (Extended Data Fig. 4a). To accomplish this, we created a library of mScarlet-I-tagged proteins, which covers almost all Rab GTPases, proteins involved in endocytosis and exocytosis, proteins related to membrane trafficking at early/recycling endosomes and Golgi/TGN, cytoskeleton-related proteins, organelle markers, and other proteins possibly related to intracellular membrane trafficking (Fig. 3a). Subsequently, we transiently expressed each of these mScarlet-I-tagged proteins in cells genome-edited for AP2-670nano^+/+^ and stably expressing the EGFP- PH(PLCδ1)-Aux1 sensor. We then performed triple-color time-lapse TIRF imaging for each protein. By implementing an imaging analysis routine to identify AP2-negative PH(PLCδ1)-Aux1-positive fusion events, we analyzed the possible association and recruitment dynamics of each of the mScarlet-I-tagged protein during the transient recruitment of the PH(PLCδ1)-Aux1 sensor at the plasma membrane (Fig. 3 and Extended Data Fig. 4a).

**Fig. 3.**
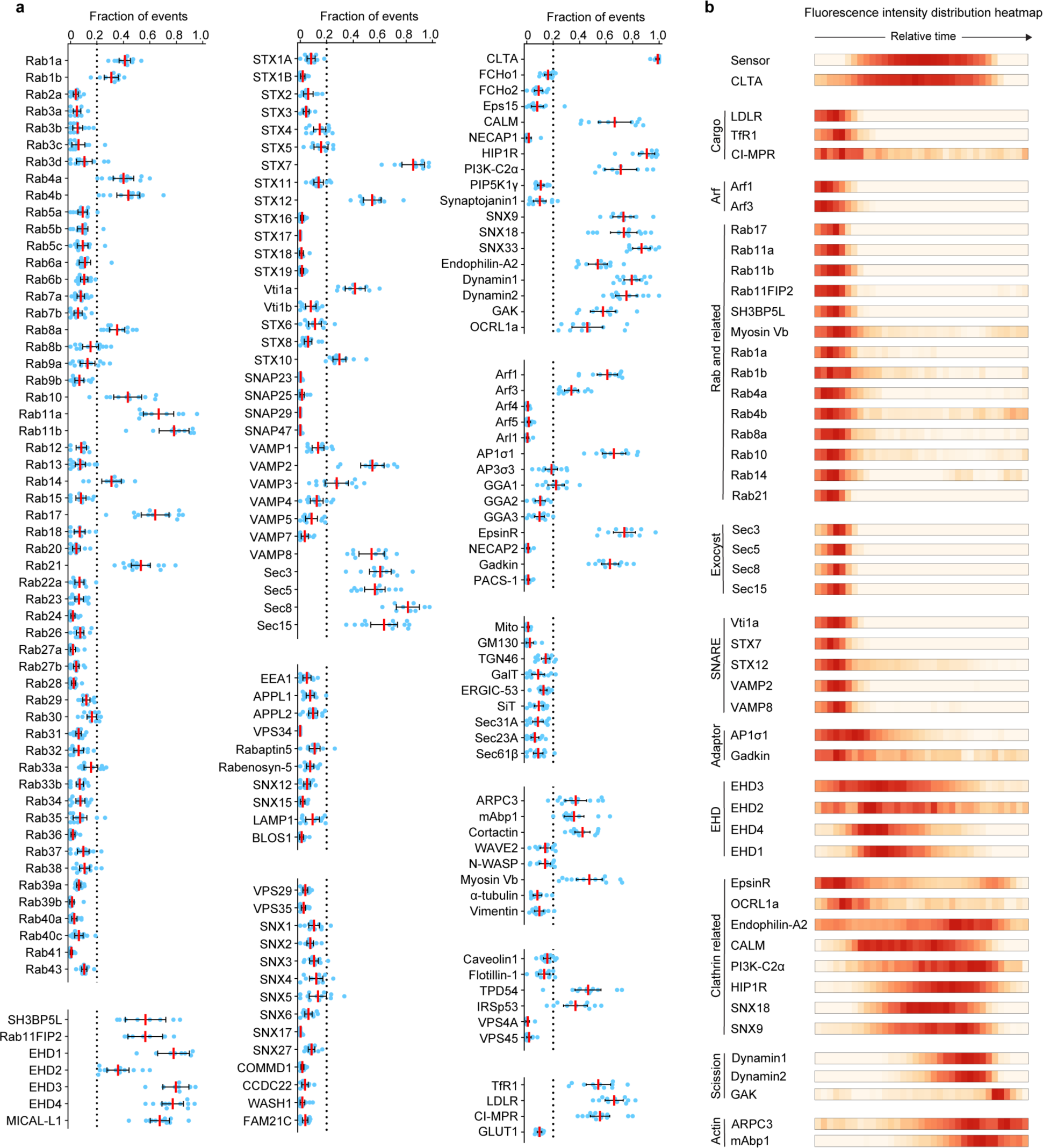
Imaging-based screening of proteins associated with AP2-negative carriers that recruited the EGFP-sensor. **a**, AP2-670nano^+/+^ cells stably expressing EGFP-sensor were transiently transfected with various proteins tagged with mScarlet-I and then imaged by TIRF microscopy at the bottom surface every 1 s for 180 s. The AP2-negative events that recruited EGFP-sensor from the time series were identified using automated 2D detection/tracking. The fraction of events containing significant signals from the mScarlet-I-tagged protein was plotted for each protein (mean ± 95% CI, n = 7-15 cells). When the averaged fraction of associated events exceeded 0.20 (dotted line), the protein was considered to be associated with the AP2-negative sensor-positive CARP carriers (see Extended Data Fig. 4d). **b**, The relative timing of the recruitment of each protein to the EGFP-sensor. The fluorescence intensity traces of the mScarlet-I-tagged related proteins were averaged and then plotted as fluorescence intensity distribution heatmaps.

We found that proteins such as Arf1/Arf3, Rab1a/Rab1b, Rab11a/Rab11b, myosin Vb, and several receptors were associated with the intracellular carriers and subsequently vanished upon sensor recruitment (Fig. 3b and Extended Data Fig. 4b). In addition to VAMP2, we observed the transient recruitment of other SNARE proteins, including VAMP8, Syntaxin 7 (STX7), STX12, and Vti1a, preceding the appearance of the sensor signal at the plasma membrane (Fig. 3b and Extended Data Fig. 4b). Furthermore, proteins such as Eps15-homology domain (EHD) proteins, HIP1R, and SNX9/SNX18, were recruited after membrane fusion and disappeared before or concurrently with the PH(PLCδ1)-Aux1 sensor. Interestingly, proteins including dynamin1/dynamin2, GAK, and actin-regulating or binding proteins were recruited during the late stages of the membrane fusion (Fig. 3b and Extended Data Fig. 4b). On the other hand, other plasma membrane-associated proteins such as flotillin-1/2, components/markers of retromers, COPII vesicles, late endosomes, and lysosomes, were not found to be related to the fusion events (Fig. 3a and Extended Data Fig. 4c). These unrelated proteins were further utilized to establish the threshold for identifying proteins with a stronger association with the recycling carriers (Extended Data Fig. 4d).

The screening results obtained from the ectopically expressed proteins were further validated by generating a series of genome-edited cell lines with endogenous proteins tagged with fluorescent proteins (see Figs 4- 6, Extended Data Fig. 4e, and Extended Data Figs 7-9). Furthermore, upon analyzing the endocytic events that were positive for both AP2 and the sensor in the screen, we observed the anticipated strong association with components of endocytic clathrin-coated pits, including clathrin, Fcho1/2, Eps15, and Eps15R, but not with known unrelated proteins such as Rab7a/Rab7b, Rab11a/Rab11b, LAMP1, and AP1/AP3 (Extended Data Fig. 5a). As further validation of the screening results obtained with the coincidence detecting sensor, we transiently expressed these related and unrelated proteins in the genome-edited cells expressing CLTA- mEGFP and AP2-670nano. Consistent with the results obtained with the sensor, we verified the recruitment or absence of each of these proteins to motile AP2-negative clathrin-positive carriers during their period of stalling at the plasma membrane (Extended Data Fig. 5b).

**Fig. 4.**
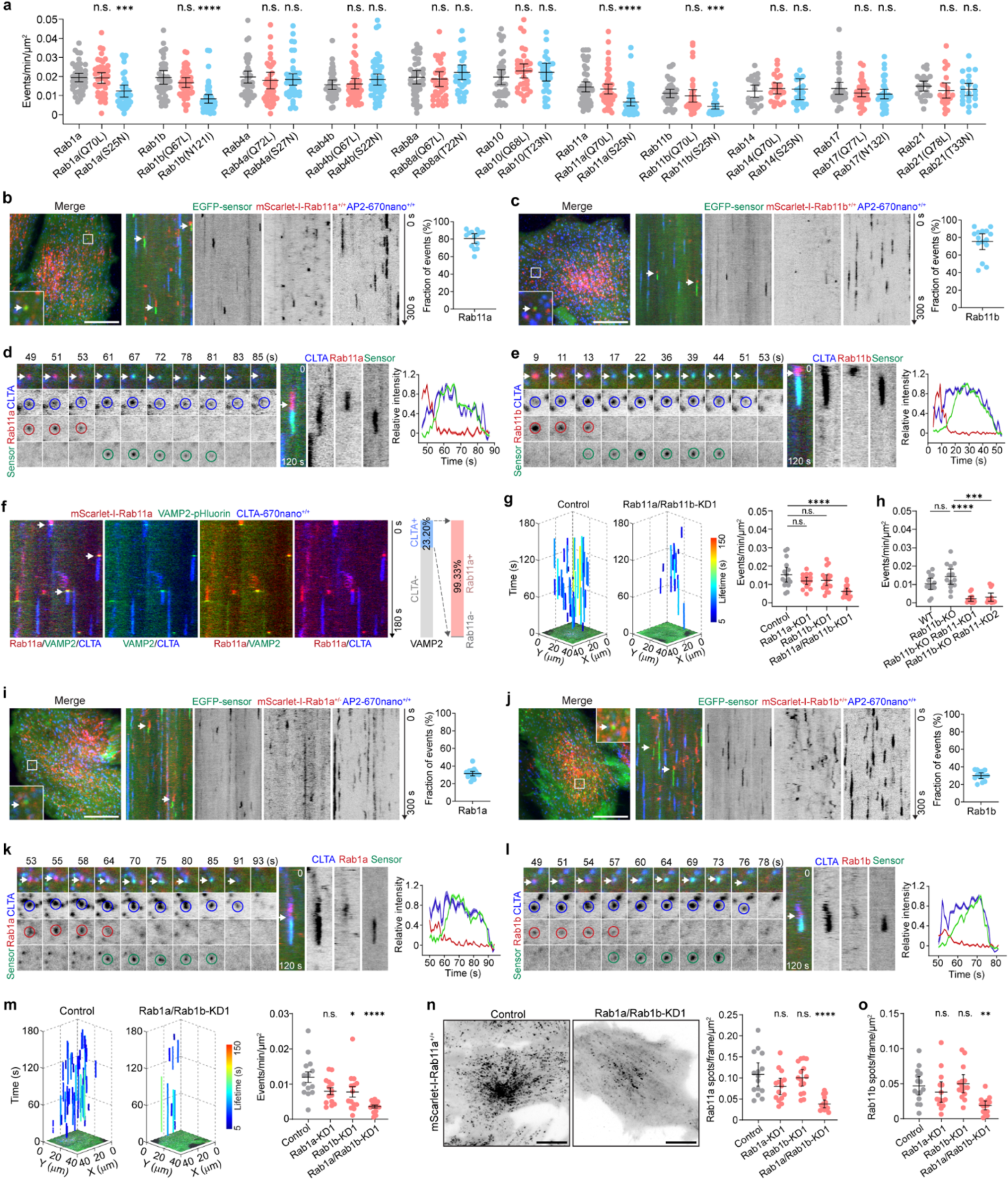
Regulation of CARP by Rab11 and Rab1. **a**, AP2-670nano^+/+^ cells stably expressing EGFP-sensor were transiently transfected with constructs expressing the indicated Rab proteins tagged with mScarlet-I, and then imaged by TIRF microscopy at the bottom surface every 1 s for 180 s. The plots show the frequency of AP2-negative sensor-positive events (fusion events) (n = 22 to 51 cells). **b**,**c**, AP2-670nano^+/+^ cells stably expressing EGFP-sensor were genome-edited to express mScarlet-I- Rab11a^+/+^ (**b**) or mScarlet-I-Rab11b^+/+^ (**c**). The T-projection and kymographs from a representative time series are shown in the left and middle panels. The Rab11a- or Rab11b-positive AP2-negative events that recruited the sensor are highlighted by arrows. The fraction of fusion events that recruited Rab11a or Rab11b is shown in the right panels (n = 14 and 15 cells). **d**,**e**, CLTA-670nano^+/+^ cells stably expressing EGFP-sensor were transiently transfected with mScarlet-I- Rab11a (**d**) or mScarlet-I-Rab11b (**e**). Representative montages, kymographs, and relative fluorescence intensity traces show the dynamic recruitment of EGFP-sensor to the clathrin/Rab11a- or clathrin/Rab11b- positive structures at the plasma membrane. **f**, CLTA-670nano^+/+^ cells stably expressing VAMP2-pHluorin were transiently transfected with mScarlet- I-Rab11a. Kymographs from a representative time series show the sequential recruitment of VAMP2 and clathrin to the Rab11a-positive exocytosis events (arrows). The fraction of VAMP2-positive exocytosis events that contained clathrin and the fraction of VAMP2/clathrin-positive events that contained Rab11a are shown on the right (2569 events from 10 cells). **g**, AP2-TagRFP^+/+^ CLTA-670nano^+/+^ cells stably expressing EGFP-sensor were treated with either control siRNA or siRNA targeting Rab11a alone, Rab11b alone, or both Rab11a and Rab11b (Rab11a/Rab11b- KD1). Left: Tracks of all fusion events from representative cells are plotted in 3D (color-coded for lifetimes). Right: Frequency of fusion events from the cells (n = 16, 16, 17, and 16 cells). **h**, Rab11b was knocked out in AP2-670nano^+/+^ mScarlet-I-Rab11b^+/+^ cells stably expressing EGFP-sensor, and then the cells were treated with control siRNA (Rab11b-KO) or two different siRNAs targeting Rab11a (KD1, KD2). The plots show the frequency of fusion events from the cells (n = 13, 13, 13, and 13 cells). **i**,**j**, AP2-670nano^+/+^ cells stably expressing EGFP-sensor were genome-edited to express mScarlet-I- Rab1a^+/-^ (**i**) or mScarlet-I-Rab1b^+/+^ (**j**). Shown are the T-projection and kymographs from representative time series and the fraction of fusion events that recruited Rab1a (n = 18 cells) or Rab1b (n = 14 cells). **k**,**l**, CLTA-670nano^+/+^ cells stably expressing EGFP-sensor were transiently transfected with mScarlet-I- Rab1a (**k**) or mScarlet-I-Rab1b (**l**). Representative montages, kymographs, and relative fluorescence intensity traces show the recruitment of EGFP-sensor to the clathrin/Rab1a- or clathrin/Rab1b-positive structures at the plasma membrane. **m**, AP2-TagRFP^+/+^ CLTA-670nano^+/+^ cells stably expressing EGFP-sensor were treated with control siRNA or siRNA targeting Rab1a alone, Rab1b alone, or both Rab1a and Rab1b. Left: Tracks of all fusion events from representative cells. Right: The frequency of fusion events from the cells (n = 14, 14, 14, and 14 cells). **n,o**, AP2-670nano^+/+^ mScarlet-I-Rab11a^+/+^ (**n**) or AP2-670nano^+/+^ mScarlet-I-Rab11b^+/+^ (**o**) cells stably expressing EGFP-sensor were treated with control siRNA or siRNA targeting Rab1a alone, Rab1b alone, or both Rab1a and Rab1b. T-projections of mScarlet-I-Rab11a from representative time series are shown in the left panels of (**n)**. The plots show the frequency of fluorescent spots of Rab11a (n = 15, 15, 15, and 15 cells) or Rab11b (n = 15, 15, 15, and 15 cells) at the plasma membrane of siRNA-treated cells. Statistical analysis was performed using the ordinary one-way ANOVA with Tukey’s multiple comparisons test; n.s., no significance; \**P* < 0.05, \*\**P* < 0.01, \*\*\**P* < 0.001, \*\*\*\**P* < 0.0001. Data are shown as mean ± 95% CI in **a**-**c**, **g**-**j**, and **m**-**o**. Cells were imaged by TIRF microscopy at the bottom surface in **a**-**n**. Scale bars, 10 μm.

In summary, our imaging-based screening and analysis successfully identified a distinct set of proteins that are recruited at different stages of intracellular clathrin-associated carrier trafficking (Fig. 3b). These proteins are likely involved in the generation, transportation, membrane fusion, or retrieval of clathrin-associated carriers.

### Regulation of CARP by Rab11 and Rab1

Rab GTPases are a family of GTPases that define membrane identity and regulate various membrane trafficking processes in mammalian cells^33^. Among the Rabs we examined, Rab11a, Rab11b, Rab17, Rab21, Rab10, Rab4a, Rab4b, Rab1a, Rab1b, Rab8a, and Rab14 exhibited association with the AP2-negative sensor-positive fusion events, mostly occurring before membrane fusion (Fig. 3). To investigate the functional relevance of these Rabs with clathrin-associated recycling carriers, we transiently expressed the above-mentioned Rabs in the mScarlet-I-tagged wild-type, dominant active, or dominant negative forms in AP2-670nano^+/+^ cells stably expressing the EGFP-PH(PLCδ1)-Aux1 sensor (Fig. 4a). We found that the dominant negative Rab11a, Rab11b, Rab1a, and Rab1b significantly decreased the frequency of AP2- negative sensor-positive fusion events (Fig. 4a). However, the dominant negative Rab4a, Rab4b, and other Rabs did not appear to affect the frequency of fusion events (Fig. 4a). These results suggest that Rab11a/Rab11b and Rab1a/Rab1b may be involved in regulating the formation or transport of clathrin-associated recycling carriers.

Rab11 is one of the key regulators of cargo recycling and is primarily enriched in perinuclear endocytic recycling compartments and recycling carriers in peripheral regions^34^. To verify the spatiotemporal and functional involvement of Rab11 in the generation or transport of clathrin-associated recycling carriers, we tagged endogenous Rab11a and Rab11b with fluorescent proteins to investigate their dynamics (Extended Data Fig. 6a,b) and eliminated the expression of Rab11a or/and Rab11b to assess their functional relevance. Live-cell imaging of genome-edited cells for Rab11a and Rab11b fused to mScarlet-I showed their accumulation on vesicles in the central area of cells, and in small highly motile vesicles near the plasma membrane (Fig. 4b,c). Tagged Rab11a and Rab11b were highly associated (∼80%) with the carriers we identified (Fig. 4b,c); they moved beneath the plasma membrane, stalled, and then recruited the PH(PLCδ1)-Aux1 sensor (Fig. 4b,c and Supplementary Videos 9 and 10). Notably, while the intracellular carriers contain both Rab11 and clathrin, Rab11a/Rab11b but not clathrin was dissociated from the carriers upon sensor recruitment (Fig. 4d,e). Furthermore, by imaging mScarlet-I-Rab11a together with VAMP2- pHluorin in the CLTA-670nano^+/+^ cells (without sensor expression), we further confirmed that nearly all the clathrin-positive exocytosis events contained Rab11a (Fig. 4f). Additionally, the Rab11 GEF molecule SH3BP5L and effector protein Rab11FIP2 were also found to associate with the vesicles before their fusion to the plasma membrane and recruitment of the PH(PLCδ1)-Aux1 sensor (Extended Data Fig. 6c).

Simultaneous elimination of Rab11a and Rab11b expression, either through dual knockdown (Extended Data Fig. 6d) or by knocking out of Rab11b followed by knockdown of Rab11a (Extended Data Fig. 6e), strongly reduced the fusion frequency of the carriers with the plasma membrane (Fig. 4g,h). Elimination of either Rab11a or Rab11b alone did not affect the fusion frequency (Fig. 4g,h), indicating the functional redundancy of Rab11a and Rab11b in regulating clathrin-associated recycling carriers. Thus, Rab11 associates with and regulates the delivery or fusion of clathrin-associated recycling carriers with the plasma membrane. Rab4 is another Rab GTPase that primarily regulates cargo recycling from early endosomes back to the plasma membrane^33^. Consistent with the results obtained from dominant negative Rab4 (Fig. 4a), we further demonstrated that the knockdown of Rab4 did not affect the fusion of the recycling carriers with the plasma membrane (Extended Data Fig. 6f,g).

The observation that constitutively negative Rab1a and Rab1b decreased the frequency of fusion events (Fig. 4a) indicates their possible involvement in regulating the formation or transportation of clathrin-associated carriers. Ectopically overexpressed Rab1a/Rab1b was mainly detected at the ER–Golgi interface and Golgi apparatus^35^. Simultaneous knockout of Rab1a and Rab1b caused cell death, while conditional knockout of Rab1a and Rab1b affected endosome distribution, delayed transferrin recycling, and disrupted the ER-to-Golgi trafficking in the cells before cell death^36^. To investigate the possible association of endogenous Rab1a/Rab1b with clathrin-associated recycling carriers, we generated genome-edited cell lines expressing mScarlet-I-tagged endogenous Rab1a or Rab1b (Extended Data Fig. 6h,i). TIRF imaging of these cell lines revealed that Rab1a and Rab1b were mainly concentrated in regions around the central areas of cells and in highly motile vesicles or tubes underneath the plasma membrane (Fig. 4i,j and Supplementary Videos 11 and 12). Endogenous Rab1a or Rab1b were found to associate with a fraction of the recycling carriers as they approached the plasma membrane and recruited the sensor (Fig. 4i,j). Similar to Rab11a/Rab11b, Rab1a/Rab1b dissociated from the clathrin-associated carriers upon membrane fusion and sensor recruitment (Fig. 4k,l). Knockdown of Rab1a and Rab1b simultaneously significantly reduced the fusion frequency of the carriers with the plasma membrane (Fig. 4m and Extended Data Fig. 6j,k). Interestingly, endogenous Rab1a and Rab1b were detected on one side of EEA1-positive early endosomes and exhibited partial association with Rab11a/Rab11b-positive vesicles (Extended Data Fig. 6l). Consistent with this observation, simultaneous knockdown of Rab1a and Rab1b expression resulted in a significant alteration in the numbers and dynamics of Rab11a/Rab11b-labeled recycling carriers beneath the plasma membrane (Fig. 4n,o and Extended Data Fig. 6m). The highly mobile vesicles or tubes observed in the control cells were replaced by relatively smaller static vesicles in the Rab1a/Rab1b-depleted cells (Fig. 4n,o and Extended Data Fig. 6m). However, the Rab11a/Rab11b-labeled recycling endosomes located around the perinuclear regions were largely unaffected (Extended Data Fig. 6n). These results suggest that Rab1 plays a crucial role in regulating the generation of Rab11-positive recycling carriers, including the clathrin-associated recycling carriers.

### Regulation of CARP by Arf1

The ADP-ribosylation factor (ARF) GTPase proteins constitute another important protein family involved in the regulation of intracellular trafficking^37^. The six mammalian ARF proteins are classified into Class I (Arf1, Arf2, and Arf3; humans lack Arf2), Class II (Arf4 and Arf5), and Class III (Arf6, localized at the plasma membrane) based on sequence homology^37^. Through the imaging-based screening, we found that Arf1 was highly related to the carriers we identified (Fig. 3). A subset of carriers also contained Arf3, while Arf4 and Arf5 showed no association (Fig. 3). To investigate the possible association of endogenous Arf1 and Arf3 with clathrin-associated carriers, we generated genome-edited cell lines expressing mScarlet-I- tagged endogenous Arf1 and Arf3 (Extended Data Fig. 7a,b). Nearly 80% of the AP2-negative sensor-positive fusion events were derived from the Arf1-positive carriers (Fig. 5a and Supplementary Video 13). Approximately one-third of these fusion events contained Arf3 (Fig. 5b). The appearance of the PH(PLCδ1)-Aux1 sensor coincided with the disappearance of Arf1 or Arf3.

**Fig. 5.**
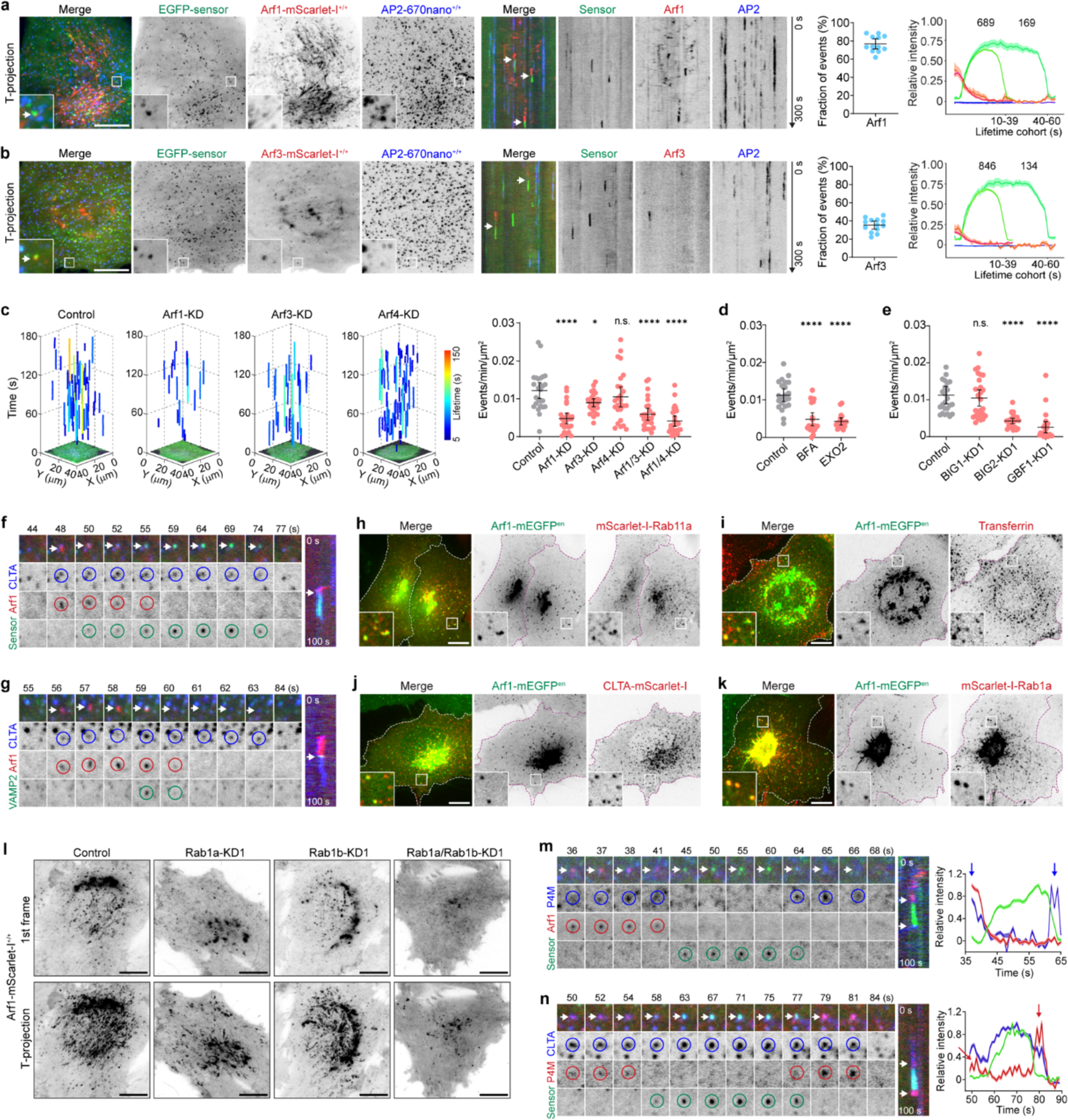
Regulation of CARP by Arf1. **a**,**b**, AP2-670nano^+/+^ cells stably expressing EGFP-sensor were genome-edited to express Arf1-mScarlet- I^+/+^ (**a**) or Arf3-mScarlet-I^+/+^ (**b**). From left to right: T-projections and kymographs from a representative time series, and the fraction and intensity cohorts of AP2-negative sensor-positive events (fusion events) that recruited Arf1 or Arf3 (n = 12 and 13 cells). Arrows in kymographs indicate Arf1 or Arf3-positive fusion events. **c**, AP2-TagRFP^+/+^ CLTA-670nano^+/+^ cells stably expressing EGFP-sensor were treated with either control siRNA or siRNA targeting Arf1 alone, Arf3 alone, Arf4 alone, Arf1 and Arf3 (Arf1/3-KD), or Arf1 and Arf4 (Arf1/4-KD). Shown are tracks of all fusion events from representative cells and the frequency of fusion events from the cells (n = 26, 26, 26, 26, 26, and 26 cells). **d**, The frequency of fusion events from AP2-TagRFP^+/+^ CLTA-670nano^+/+^ cells stably expressing EGFP- sensor and treated with DMSO (control), BFA, or EXO2 (n = 22, 21, and 19 cells). **e,** AP2-TagRFP^+/+^ CLTA-670nano^+/+^ cells stably expressing EGFP-sensor were treated with either control siRNA or siRNA targeting BIG1, BIG2, or GBF1. The plots show the frequency of fusion events from the cells (n = 25, 26, 25, and 25 cells). **f**, CLTA-670nano^+/+^ cells stably expressing EGFP-sensor were transiently transfected with Arf1-mScarlet-I. The representative montage and kymograph show the recruitment of EGFP-sensor to the clathrin- and Arf1-containing carrier (arrows) at the plasma membrane. **g**, CLTA-670nano^+/+^ cells were transiently transfected with VAMP2-pHluorin and Arf1-mScarlet-I. The representative montage and kymograph show the burst recruitment of VAMP2-pHluorin to an Arf1- and clathrin-associated carrier (arrows) at the plasma membrane. **h-k**, Genome-edited Arf1-mEGFP cells (pool) were transiently transfected with mScarlet-I-Rab11a (**h**), CLTA-mScarlet-I (**j**), or mScarlet-I-Rab1a (**k**), or incubated with Alexa Fluor 568-conjugated transferrin for 5 min (**i**), and then imaged by spinning-disk confocal microscopy at the plane near the bottom surface. **l**, AP2-670nano^+/+^ Arf1-mScarlet-I^+/+^ cells stably expressing EGFP-sensor were treated with either control siRNA or siRNA targeting Rab1a alone, Rab1b alone, or both Rab1a and Rab1b (Rab1a/Rab1b-KD1). Shown are the first frame and the T-projection of Arf1-mScarlet-I from representative time series. **m**, Cells stably expressing EGFP-sensor were transiently transfected with Arf1-mScarlet-I and the clathrin- specific PI4P sensor Halo-P4M(DrrA)-Aux1. The representative montage, kymograph, and relative fluorescence intensity traces show sequential burst recruitments (arrows) of Halo-P4M(DrrA)-Aux1 during the initial and final stages of EGFP-sensor recruitment to an Arf1-positive carrier at the plasma membrane. **n**, CLTA-670nano^+/+^ cells stably expressing EGFP-sensor were transiently transfected with mScarlet-I- P4M(DrrA)-Aux1. The representative montage, kymograph, and relative fluorescence intensity traces show sequential burst recruitments (arrows) of mScarlet-I-P4M(DrrA)-Aux1 during the initial and final stages of EGFP-sensor recruitment to a clathrin-positive carrier at the plasma membrane. Statistical analysis was performed using the ordinary one-way ANOVA with Tukey’s multiple comparisons test; n.s., no significance; \**P* < 0.05, \*\*\*\**P* < 0.0001. Data are shown as mean ± 95% CI in **a-e**; mean ± SEM in cohorts in **a** and **b**. Cells were imaged by TIRF microscopy at the bottom surface in **a-g** and **l-n**. Scale bars, 10 μm.

To determine the functional relevance of Arf1 and Arf3, we reduced the expression of Arf1, Arf3, and Arf4 (as a control). Consistent with the imaging results, the knockdown of Arf1, Arf3, or Arf4 strongly, slightly, or minimally reduced the fusion frequency of the carriers with the plasma membrane, respectively (Fig. 5c and Extended Data Fig. 7c). We observed that Arf1 knockdown did not affect the expression of Arf3, whereas Arf3 knockdown resulted in an upregulation of Arf1 expression (Extended Data Fig. 7c). Furthermore, we confirmed that knockout of Arf1 expression in genome-edited Arf1-mScarlet-I^+/+^ cells reduced the fusion frequency, and the reduction could be rescued by re-expression of the wild-type or constitutively active form of Arf1, but not the dominant negative form (Extended Data Fig. 7d). The activation and membrane recruitment of Arf1 is mediated by Arf-GEFs (GBF1, BIG1, and BIG2), which can be inactivated by brefeldin A (BFA). BFA treatment induced a dispersed distribution of endogenous Arf1 and reduced the fusion frequency (Fig. 5d). EXO2, a BFA-like inhibitor, exhibited a similar inhibitory effect (Fig. 5d). Accordingly, knockdown of GBF1 or BIG2 expression effectively reduced the frequency of fusion events (Fig. 5e and Extended Data Fig. 7e,f). These findings imply that Arf1 is involved in the regulation of clathrin-associated recycling carriers. Indeed, Arf1 is associated with the intracellular clathrin-associated carriers before recruitment of the PH(PLCδ1)-Aux1 sensor (Fig. 5f). Furthermore, in the absence of the sensor expression, the fusion of intracellular Arf1/clathrin-positive carriers with the plasma membrane could still be detected through the burst signal of VAMP2-pHluorin (Fig. 5g and Extended Data Fig. 7g). Over 80% of clathrin-positive exocytosis events contained Arf1 (Extended Data Fig. 7g).

Live-cell imaging of genome-edited cells revealed that Arf1 is mainly concentrated in Golgi regions and enriched in highly motile tubes or vesicles beneath the plasma membrane (Fig. 5h-k). Notably, these Arf1- positive tubes or vesicles near the plasma membrane were also positive for Rab11a/Rab11b (Fig. 5h and Extended Data Fig. 7h) as well as the fluorescently labeled transferrin 5 min after their internalization (Fig. 5i). This observation suggests that these tubulovesicular carriers are likely involved in the sorting or recycling of endocytosed cargoes. As expected, these Arf1-positive tubulovesicular carriers near the plasma membrane were positive for clathrin (Fig. 5j). By conducting live-cell tracking at a high imaging rate to capture these intracellular highly motile Arf1-positive tubular structures before their fusion and sensor recruitment at the plasma membrane, we found that clathrin signal was typically observed at the rear part of these highly motile tubules (Extended Data Fig. 7j). Upon the cessation of movement near the plasma membrane, these clathrin-containing tubular Arf1 structures transitioned into vesicular structures, coinciding with sensor recruitment (Extended Data Fig. 7j). Live-cell SIM imaging further showed that clathrin localizes at subregions of these tubular Arf1 carriers (Extended Data Fig. 7k).

Interestingly, the tubes or vesicles labeled with Arf1 also contain Rab1a and Rab1b (Fig. 5k and Extended Data Fig. 7i). Simultaneous knockdown of Rab1a and Rab1b expression strongly decreased the presence of Arf1-positive tubes or vesicles near the plasma membrane (Fig. 5l). However, the knockdown of Arf1 did not affect the intracellular distribution of endogenous Rab1a/Rab1b or Rab11a/Rab11b (Extended Data Fig. 7l), Nonetheless, it led to a decrease in the distribution of AP1 in the perinuclear regions (Extended Data Fig. 7l). Together, these findings show that Rab1 is crucial for the generation of Arf1/Rab11/clathrin-positive carriers beneath the plasma membrane.

The subcellular localization of Arf1 is closely related to the generation of phosphatidylinositol-4-phosphate (PI4P)^38^. The recruitment of adaptor proteins, such as AP1, to intracellular membranes requires PI4P^39^. PI4P regulates vesicular transport at TGN and cargo sorting at early endosomes^39, 40^. To investigate the potential involvement of PI4P in clathrin-associated recycling carriers, we expressed the clathrin-coated structure-specific PI4P sensor Halo-P4M(DrrA)-Aux1^11^ in cells stably expressing the EGFP-PH(PLCδ1)- Aux1 sensor. Along with Arf1 and clathrin, the PI4P sensor was recruited to recycling carriers before their fusion with the plasma membrane (Fig. 5m,n). Intriguingly, a strong signal from the PI4P sensor was observed after the disappearance of the PH(PLCδ1)-Aux1 sensor (Fig. 5m,n), indicating local generation of PI4P in recycling carriers following their detachment from the plasma membrane. Indeed, OCRL, a 5-phosphatase that converts PI(4,5)P_2_ to PI4P, was similarly recruited to recycling carriers after the disappearance of the PH(PLCδ1)-Aux1 sensor signal (Extended Data Fig. 7m). The enrichment of PI4P in clathrin-associated carriers before membrane fusion and after budding was further supported by the similar recruitment of endogenous EpsinR (Extended Data Fig. 7n), a protein that binds to PI4P in clathrin/AP1- containing intracellular vesicles^41^.

### The partially fused clathrin-associated recycling carriers are released from the plasma membrane by dynamin

Dynamin is a GTPase that catalyzes membrane fission at the neck of a budding vesicle upon GTP hydrolysis^42^. An unexpected observation was that ectopically overexpressed dynamin was recruited to AP2- negative carriers after membrane fusion (Fig. 3 and Extended Data Fig. 8a). We also observed that after staying at the plasma membrane for a few to tens of seconds, the membrane-fused clathrin-associated recycling carriers moved away from the fusion site and lost the sensor signal (Fig. 1h-k). These observations led us to speculate that the carriers partially fused with the plasma membrane and then reentered the cells after dynamin-mediated scission. To validate this hypothesis, we generated a genome-edited cell line expressing mScarlet-I-tagged endogenous dynamin2 (Extended Data Fig. 8b), which is the ubiquitously expressed dynamin isoform in mammals^42^. TIRF imaging of the genome-edited cells confirmed that, apart from the typical AP2/clathrin-coated endocytic vesicles, dynamin2 was recruited to the AP2-negative PH(PLCδ1)-Aux1 sensor-positive fusion events (Fig. 6a,b and Supplementary Video 14). Approximately 90% of the fusion events showed recruitment of endogenous dynamin2 during the late stage of fusion (Fig. 6c). Compared to dynamin2 recruited to the AP2-positive endocytic vesicles, dynamin2 recruited to the AP2-negative fusion events exhibited similar dynamics but lower fluorescence intensity (Fig. 6d,e). We further validated the recruitment of dynamin2 to fusion events derived from Arf1-, AP1-, Rab1a, or Rab11a- containing recycling carriers (Fig. 6f-i). The recruitment of dynamin2 to the partially fused carriers was also confirmed in two additional cell lines (Extended Data Fig. 8c,d). Additionally, endophilin-A2, which is closely related to the recruitment and activation of dynamin, displayed similar recruitment dynamics as dynamin to clathrin-associated carriers (Extended Data Fig. 8e).

**Fig. 6.**
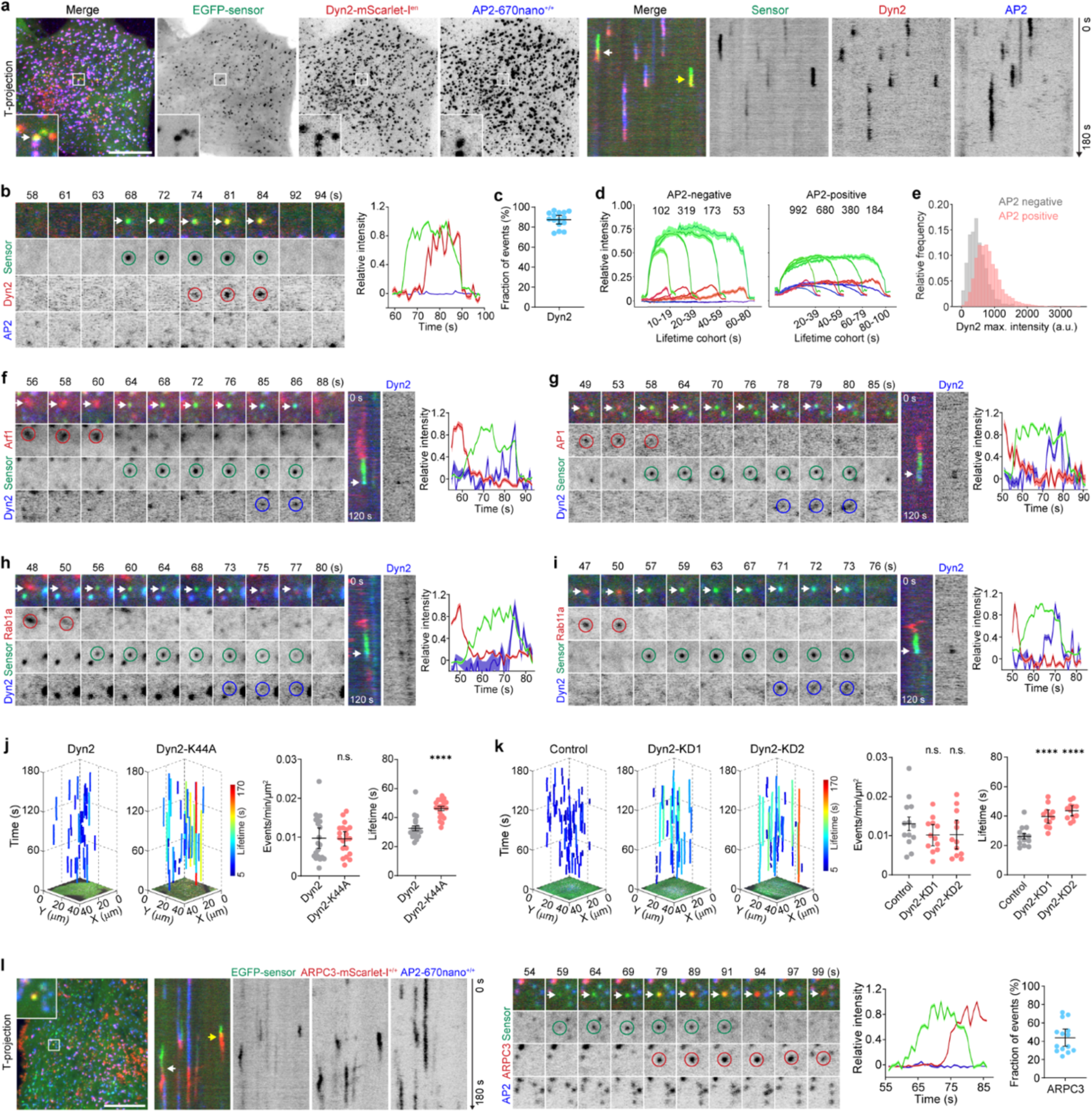
**Clathrin-associated recycling carriers are released from the plasma membrane by dynamin. a-e**, AP2-670nano^+/+^ cells stably expressing EGFP-sensor were genome-edited to express dynamin2- mScarlet-I (pool, Dyn2-mScarlet-I^en^). T-projections and kymographs are shown from a representative time series (**a**). Representative montage and relative fluorescence intensity traces show the recruitment of dynamin2 during the late stage of sensor recruitment to an AP2-negative carrier (arrows) (**b**). The fraction of AP2-negative sensor-positive fusion events that recruited dynamin2 (**c**). The intensity cohorts of AP2- negative fusion events (left) or AP2-positive endocytic events (right) that recruited dynamin2 (**d**). The distribution of the max. fluorescence intensity of dynamin2 from the AP2-negative or AP2-positive events (n = 15 cells) (**e**). **f-i**, Cells stably expressing EGFP-sensor were transiently transfected with Dyn2-670nano together with Arf1-mScarlet-I (**f**), AP1σ1-mScarlet-I (**g**), mScarlet-I-Rab1a (**h**), or mScarlet-I-Rab11a (**i**). Representative montages, kymographs, and relative fluorescence intensity traces show the recruitment of dynamin2 (arrows) during the late stage of sensor recruitment to the Arf1-, AP1-, Rab1a-, or Rab11a-positive carrier. **j**, AP2-670nano^+/+^ cells stably expressing EGFP-sensor were transiently transfected with Dyn2-mScarlet-I or Dyn2-K44A-mScarlet-I. Shown are tracks of all fusion events from representative cells, and the frequency and lifetime of fusion events from the cells (n = 20 and 20 cells). **k**, AP2-TagRFP^+/+^ CLTA-670nano^+/+^ cells stably expressing EGFP-sensor were treated with control siRNA or two different siRNAs targeting dynamin2. Shown are tracks of all fusion events from representative cells, and the frequency and lifetime of fusion events from the cells (n = 13, 13, and 12 cells). **l**, AP2-670nano^+/+^ cells stably expressing EGFP-sensor were genome-edited to express ARPC3-mScarlet- I^+/+^. From left to right: the T-projection and kymographs from a representative time series; the montage and relative fluorescence intensity traces of the event indicated by the yellow arrow in the kymograph; and the fraction of fusion events that recruited ARPC3 (n = 14 cells). Statistical analysis was performed using the unpaired, two-tailed Student’s *t*-test in **j**, and the ordinary one- way ANOVA with Tukey’s multiple comparisons test in **k**; n.s., no significance; \*\*\*\**P* < 0.0001. Data are shown as mean ± 95% CI in **c** and **j-l**; mean ± SEM in **d**. Cells were imaged by TIRF microscopy at the bottom surface in **a-l**. Scale bars, 10 μm.

To examine the impact of dynamin on the membrane dynamics of clathrin-associated carriers, we interfered with dynamin function by overexpressing the dominant negative dynamin2 (K44A) (Fig. 6j), reducing dynamin2 expression with two different siRNA sequences^43^ (Fig. 6k and Extended Data Fig. 8f), or treating cells with the dynamin inhibitor Dynole 2-24 (Extended Data Fig. 8g). All these interventions prolonged the association of the clathrin-associated carriers with the plasma membrane, indicating impaired scission and thus retention of recycling structures at the plasma membrane.

Actin assembly facilitates the invagination, constriction, or scission of endocytic vesicles at the plasma membrane^44^. During the imaging-based screening, we observed the recruitment of several proteins related to actin-based trafficking or actin assembly at various stages of clathrin-associated carrier formation (Fig. 3). Similar to dynamin2, ARPC3, a subunit of the Arp2/3 complex that nucleates actin filament assembly, was recruited to the carriers at the late stage of fusion (Fig. 6l and Extended Data Fig. 8h). Remarkably, the fluorescence signal of endogenous ARPC3 remains after the disappearance of the PH(PLCδ1)-Aux1 sensor signal, indicating the continuous association of ARPC3 with the carriers after dynamin-mediated fission. Continuous association with the released carriers was also observed with another actin-related protein, mAbp1 (Extended Data Fig. 8i,j). These observations further validate that instead of fully collapsing into the plasma membrane, the clathrin-associated recycling carriers undergo partial membrane fusion and are subsequently retrieved and released from the plasma membrane.

### Distinct regulation of CARP by actin and microtube cytoskeletons

The generation of transport carriers and the delivery of these carriers from intracellular compartments to the plasma membrane are facilitated by cytoskeletons and associated motors^45^. Myosin Vb, a Class V myosin responsible for actin-based vesicular trafficking of Rab11-positive vesicles^46, 47^, associated with the intracellular clathrin-associated carriers before their fusion with the plasma membrane (Extended Data Fig. 9a). The fusion frequency of clathrin-associated recycling carriers with the plasma membrane significantly decreased upon disruption of the actin cytoskeleton using inhibitors such as latrunculin A or blebbistatin (Fig. 7a,b), suggesting that the actin cytoskeleton is involved in the delivery of the intracellular carriers to the plasma membrane.

**Fig. 7.**
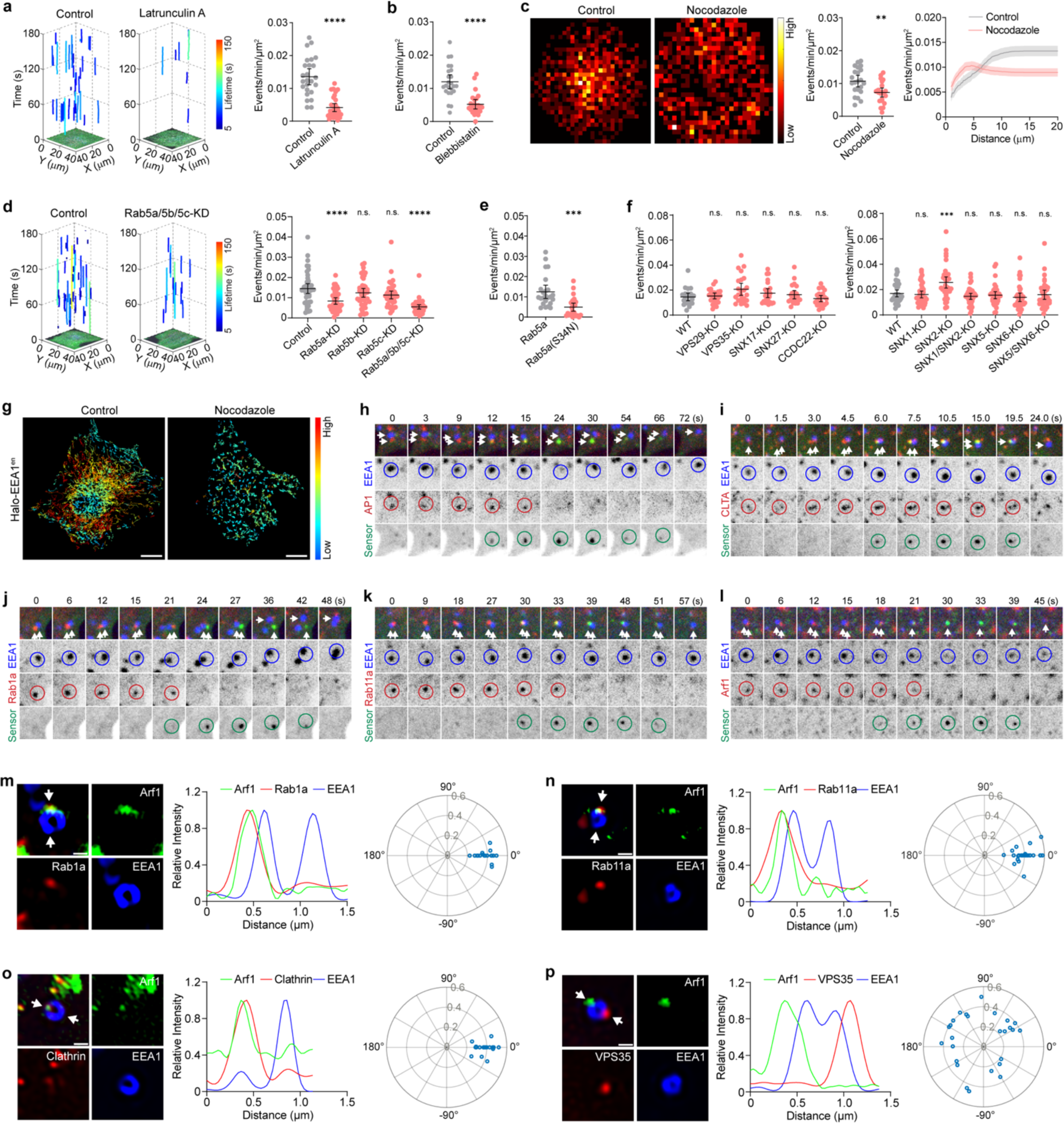
Clathrin-associated recycling carriers are derived from early endosomes. **a**, AP2-TagRFP^+/+^ CLTA-670nano^+/+^ cells stably expressing EGFP-sensor were treated with ethanol (control) or latrunculin A, and then imaged at 1.5-s intervals by TIRF microscopy. Shown are tracks of AP2-negative sensor-positive fusion events from representative cells, and the frequency of fusion events from the cells (n = 25 and 30 cells). **b**, AP2-TagRFP^+/+^ CLTA-670nano^+/+^ cells stably expressing EGFP-sensor were treated with DMSO (control) or the myosin II inhibitor blebbistatin, and then imaged by TIRF microscopy. The frequency of fusion events from the cells is shown (n = 24 and 24 cells). **c**, AP2-TagRFP^+/+^ CLTA-670nano^+/+^ cells stably expressing EGFP-sensor were treated with DMSO (control) or nocodazole, and then imaged at 1.5-s intervals by TIRF microscopy. Left: Spatial frequency distribution of fusion events from DMSO- or nocodazole-treated cells (n = 22 and 23 cells). Middle: Frequency of fusion events. Right: Relative frequency of fusion events occurring from the cell edge (set as 0) to the center. **d**, AP2-TagRFP^+/+^ CLTA-670nano^+/+^ cells stably expressing EGFP-sensor were treated with either control siRNA or siRNA targeting Rab5a, Rab5b, Rab5c, or Rab5a/5b/5c, and then imaged at 1-s intervals by TIRF microscopy. Tracks of all fusion events from representative cells and the frequency of fusion events from the siRNA-treated cells are shown (from left to right, n = 43, 40, 41, 41, and 41 cells). **e**, AP2-670nano^+/+^ cells stably expressing EGFP-sensor were transiently transfected with mScarlet-I-Rab5a or mScarlet-I-Rab5a(S34N), and then imaged at 1-s intervals by TIRF microscopy. The plots show the frequency of fusion events from the cells (n = 24 and 24 cells). **f**, AP2-mScarlet^+/+^ cells were subjected to knock out of the expression of the indicated proteins and then transiently transfected with EGFP-sensor, and then imaged at 1-s intervals by TIRF microscopy. The plots show the frequency of fusion events from the cells (from left to right, n = 22, 23, 23, 24, 22, 22, 36, 38, 36, 37, 36, 36, and 36 cells). **g**, Cells stably expressing EGFP-sensor and genome-edited to express Halo-EEA1 (pool) were pretreated with DMSO (control) or nocodazole for 1 h and then imaged at 1.5-s intervals by spinning-disk confocal microscopy near the bottom surface. The EEA1-positive endosomes in the time series were detected and tracked. The overlay of all tracks (color-coded based on the velocity of endosome movement) from representative cells treated with DMSO or nocodazole is shown. **h-l**, Cells stably expressing EGFP-sensor and genome-edited to express Halo-EEA1 (pool) were transiently transfected with AP1σ1-mScarlet-I (**h**), CLTA-mScarlet-I (**i**), mScarlet-I-Rab1a (**j**), mScarlet-I-Rab11a (**k**), or Arf1-mScarlet-I (**l**), stained with the JFX_650_-HaloTag ligand, treated with nocodazole for 1 h, and then imaged at 1.5-s intervals by spinning-disk confocal microscopy near the bottom surface. Montages show the recruitment of EGFP-sensor to the AP1/CLTA/Rab1a/Rab11a/Arf1-positive structures budded from the EEA1-positive endosomes close to the plasma membrane (arrows). **m-p**, Genome-edited cells expressing Arf1-mEGFP and Halo-EEA1 (pool) were transiently transfected with mScarlet-I-Rab1a (**m**), mScarlet-I-Rab11a (**n**), CLTA-mScarlet-I (**o**), or VPS35-mScarlet-I (**p**), and then imaged near the middle plane by SIM. Left: Sub-organelle distribution of Arf1 with the indicated protein on an EEA1-positive early endosome. Middle: Intensity profiles of the proteins (between the arrows) on the early endosome. Right: Polar plots of the relative spatial distribution of Rab1a, Rab11a, clathrin, or VPS35 with respect to Arf1 on individual early endosomes. Statistical analysis was performed using the unpaired, two-tailed Student’s *t*-test in **a**-**c** and **e**, and the ordinary one-way ANOVA with Tukey’s multiple comparisons test in **d** and **f**; n.s., no significance; \*\**P* < 0.01, \*\*\**P* < 0.001, \*\*\*\**P* < 0.0001. Data are shown as mean ± 95% CI in **a**-**f**. Scale bars, 10 μm in **g** and 0.5 μm in **m**-**p**.

To examine the role of microtubules in regulating clathrin-associated recycling carriers, we disrupted microtubules using nocodazole, resulting in a slight decrease in the overall frequency of fusion events (Fig. 7c). However, unlike control cells where membrane fusion primarily occurred near the central area of the plasma membrane, fusion events in cells with disrupted microtubules showed a more dispersed distribution around the periphery of the plasma membrane (Fig. 7c). This observation suggests that microtubules do not play a major role in transporting the recycling carriers to the plasma membrane but may influence the subcellular regions where membrane fusion is most likely to occur.

### Clathrin-associated recycling carriers are derived from early endosomes

The above findings have demonstrated the existence of intracellular clathrin-associated recycling carriers that transiently fuse with the plasma membrane and then return to the cells. The next question was where these carriers originated from. Previous studies and our results have shown that Arf1 is primarily concentrated in regions containing the Golgi apparatus and is also localized to transferrin-positive early endosomes^48^ (Fig. 5i). Since we did not observe a strong association of clathrin-associated recycling carriers with post-Golgi vesicles (Extended Data Fig. 3c-f), we subsequently investigated whether they are generated from early endosomes. To explore this, we interfered with the expression or function of Rab5, the key regulator of early endosome biogenesis and functions^33^. We found that the simultaneous elimination of the three isoforms of Rab5 (Rab5a, Rab5b, and Rab5c) resulted in reduced fusion frequency of the recycling carriers with the plasma membrane (Fig. 7d and Extended Data Fig. 9b). Similar results were observed by interfering with Rab5 function through overexpression of the dominant negative mutant of Rab5a (Fig. 7e). Previous studies have demonstrated that endosomal retrieval and recycling of cargoes require various endosomal retrieval complexes such as retriever, retromer, and the COMMD/CCDC22/CCDC93 (CCC) complex^47, 49^. To investigate whether the clathrin-associated recycling carriers are generated by these retrieval complexes from early endosomes, we knocked down the expression of essential subunits of SNX27-retromer, SNX3-retromer, SNX17 retriever, SNX1/2:SNX5/6, and CCC complex (Extended Data Fig. 9c,d). Unexpectedly, the elimination of these essential subunits did not decrease the frequency of fusion events (Extended Data Fig. 9c,d). This finding was further confirmed in genome-edited cells with these essential subunits knocked out (Fig. 7f and Extended Data Fig. 9e,f). Thus, the above results indicate that clathrin-associated recycling carriers are derived from early endosomes independently of the known endosomal retrieval complexes.

By imaging the genome-edited cells expressing Halo-EEA1 at the basal plane using spinning-disk confocal microscopy, we occasionally observed the transient appearance of the PH(PLCδ1)-Aux1 sensor near the motile EEA1-positive endosomes beneath the plasma membrane (Extended Data Fig. 9h). Since EEA1- positive early endosomes constantly move on microtubules and are primarily located around the perinuclear region^50^, it is technically challenging to track the generation and subsequent fusion of recycling carriers from these motile early endosomes. To overcome this limitation, we treated the cells with nocodazole to disrupt microtubules. In line with our observation that microtubule disruption led to a more dispersed distribution of fusion sites at the plasma membrane (Fig. 7c), early endosomes showed reduced movement and a more dispersed distribution near the plasma membrane in cells with disrupted microtubules^50^ (Fig. 7g and Extended Data Fig. 9g). This enables us to simultaneously capture the generation of recycling carriers from early endosomes and subsequently the recruitment of the PH(PLCδ1)-Aux1 sensor to these generated carriers. Indeed, we readily observed the recruitment of the PH(PLCδ1)-Aux1 sensor to the AP1-, CLTA-, Rab1-, Rab11-, or Arf1-containing carriers originating from the less motile early endosomes close to the plasma membrane (Fig. 7h-l, Extended Data Fig. 9i, and Supplementary Video 15). Remarkably, these protein-enriched domains were usually derived from one side of early endosomes and then recruited the PH(PLCδ1)-Aux1 sensor (Fig. 7h-l and Extended Data Fig. 9i).

Since the knockout or knockdown of endosomal retrieval complexes had no effect on the generation of clathrin-associated recycling carriers (Fig. 7f and Extended Data Fig. 9c,d), and AP1/Rab1/Rab11/Arf1 was concentrated on one side of early endosomes (Fig. 7h-l), we speculated that the AP1/Rab1/Rab11/Arf1- positive clathrin-associated recycling carriers might be generated from a distinct subregion of early endosomes. To determine the sub-organelle distribution of different components on early endosomes, we conducted super-resolution SIM imaging. We found that components of the identified recycling structures spatially colocalized with each other at one side of early endosomes (Fig. 7m-o and Extended Data Fig. 9j,k), whereas the retromer component was in a separate subregion of early endosomes (Fig. 7p). Together, these findings suggest that clathrin-associated recycling carriers are generated from a subdomain of early endosomes lacking the retrieval complexes.

### CARP mediates the fast recycling of receptors to the plasma membrane

Early endosomes are the sorting hub where internalized cargoes, including receptors, are sorted for recycling or degradation^6^. During our imaging-based screening, we found that receptors such as TfR and LDLR, which undergo constitutive recycling following clathrin-mediated endocytosis, were actively transported to the plasma membrane with clathrin-associated recycling carriers (Fig. 3 and Extended Data Fig. 4b). By imaging the membrane fusion of pHluorin-tagged TfR1, we further confirmed the recruitment of the sensor following a burst signal of TfR1-pHluorin at the plasma membrane (Fig. 8a and Extended Data Fig. 10a). We also observed the partial release of TfR1-pHluorin after membrane fusion (Fig. 8a). These results provide additional evidence supporting the partial fusion of clathrin-associated recycling carriers with the plasma membrane.

**Fig. 8.**
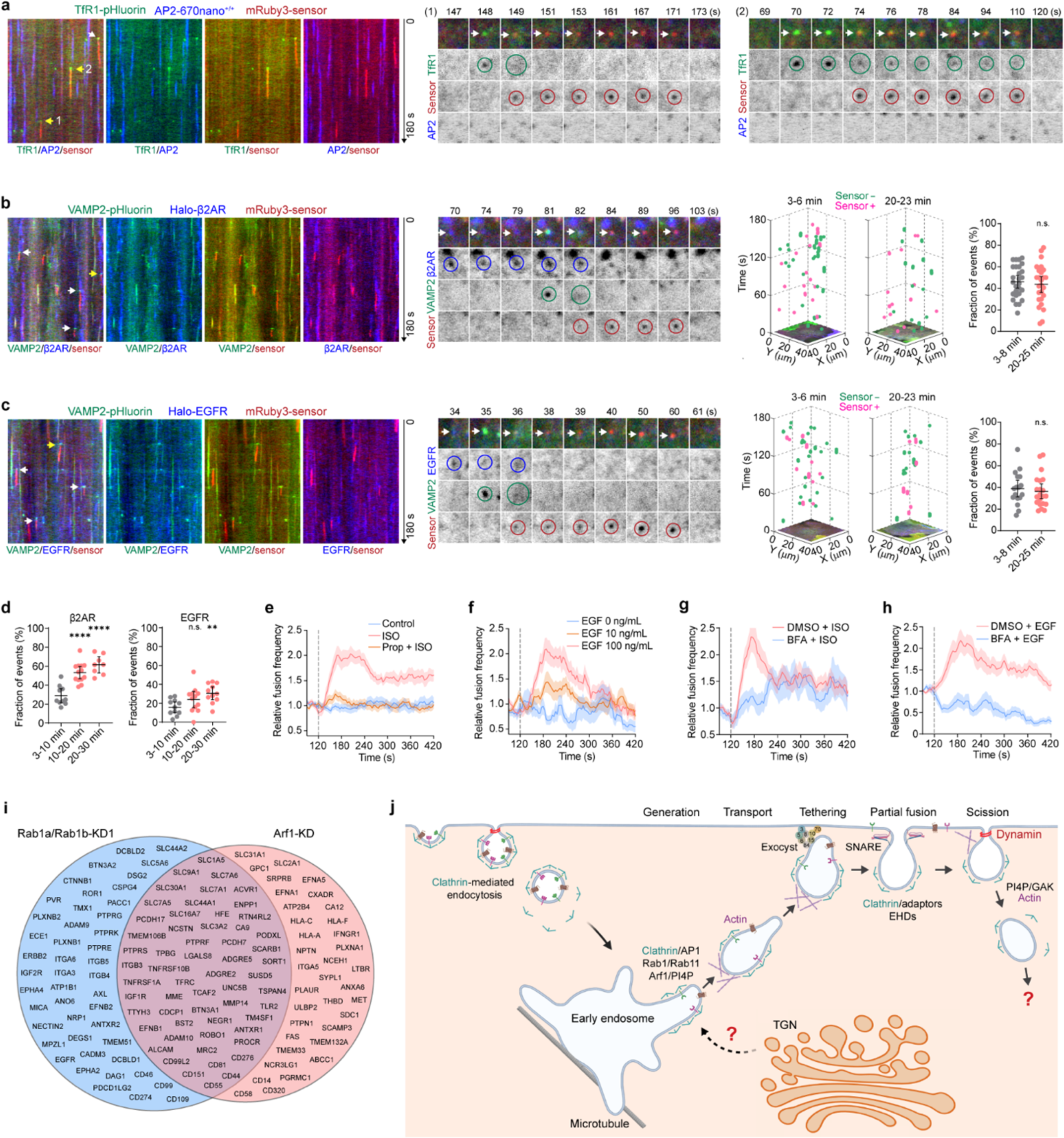
**Clathrin-associated carriers mediate the recycling of receptors back to the plasma membrane. a**, AP2-670nano^+/+^ cells stably expressing mRuby3-sensor were transiently transfected with TfR1-pHluorin, and then imaged by TIRF microscopy every 1 s for 180 s. Kymographs from a representative time series are shown on the left. Montages of the two events indicated by the yellow arrows in the kymograph show sensor recruitment to TfR1 exocytosis events. **b**, Cells stably expressing VAMP2-pHluorin and mRuby3-sensor were transiently transfected with Halo- β2AR, stained with membrane-impermeable JF_635_i-HaloTag ligand, treated with ISO for 3 min, and then imaged at 1-s intervals by TIRF microscopy. From left to right: kymographs from a representative time series; the montage showing sensor recruitment after a burst of VAMP2 during a β2AR exocytosis event; tracks of identified β2AR-VAMP2 exocytosis events without or with sensor recruitment, from representative cells during the early and late stages of ISO stimulation; and the fraction of β2AR-VAMP2 exocytosis events with sensor recruitment (n = 27 and 26 cells). **c**, Cells stably expressing VAMP2-pHluorin and mRuby3-sensor were transiently transfected with Halo- EGFR, stained with the JF_635_i-HaloTag ligand, treated with EGF for 3 min, and then imaged at 1-s intervals by TIRF microscopy. From left to right: kymographs from a representative time series; the montage showing sensor recruitment after a burst of VAMP2 during an EGFR exocytosis event; tracks of all the identified EGFR-VAMP2 exocytosis events without or with sensor recruitment, from representative cells during the early and late stages of EGF stimulation; and the fraction of EGFR-VAMP2 exocytosis events with sensor recruitment (n = 17 and 21 cells). **d**, The fraction of AP2-negative sensor-positive fusion events with β2AR or EGFR recruitment at different periods (3-10 min, 10-20 min, and 20-30 min) after ISO or EGF stimulation (β2AR: n = 10, 13, and 8 cells; EGFR: n = 11, 12, and 11 cells). **e**, AP2-TagRFP^+/+^ cells stably expressing EGFP-sensor and SNAP-β2AR were treated without or with ISO, or pretreated with the β-blocker propranolol and then ISO (Prop + ISO), at 120 s during continuous imaging at 4-s intervals by TIRF microscopy. The relative frequency of fusion events at different times is plotted (n = 32, 32, and 34 cells). **f**, AP2-TagRFP^+/+^ CLTA-670nano^+/+^ cells stably expressing EGFP-sensor were treated with different concentrations of EGF at 120 s during continuous imaging at 4-s intervals by TIRF microscopy. The relative frequency of fusion events at different times is plotted (n = 27, 73, and 65 cells). **g**, AP2-TagRFP^+/+^ cells stably expressing EGFP-sensor and SNAP-β2AR were pretreated with DMSO or BFA for 30 min, and then treated with ISO at 120 s during continuous imaging at 4-s intervals by TIRF microscopy. The relative frequency of fusion events at different times is plotted (n = 15 and 15 cells). **h**, AP2-TagRFP^+/+^ CLTA-670nano^+/+^ cells stably expressing EGFP-sensor were pretreated with DMSO or BFA for 30 min, and then treated with EGF (100 ng/ml) at 120 s during continuous imaging at 4-s intervals by TIRF microscopy. The relative frequency of fusion events at different times is plotted (n = 24 and 28 cells). **i**, Cells were treated with control siRNA or siRNA targeting Rab1a/Rab1b or Arf1. The biotinylated cell surface proteins were analyzed by mass spectrometry and compared with those of the control cells. Shown are proteins reduced more than 1.2-fold from the surfaces of Rab1a/Rab1b- or Arf1-depleted cells from three independent experiments. **j**, A model summarizing the different trafficking steps of CARP. Statistical analysis was performed using the unpaired, two-tailed Student’s *t*-test in **b** and **c**, and the ordinary one-way ANOVA with Tukey’s multiple comparisons test in **d**; n.s., no significance; \*\**P* < 0.01; \*\*\*\**P* < 0.0001. Data are shown as mean ± 95% CI in **b**-**d**; mean ± SEM in **e**-**h**.

To further explore whether clathrin-associated carriers are functionally related to the trafficking of signaling receptors, we tracked the endocytic recycling of β2AR from the GPCR families and the epidermal growth factor receptor (EGFR) from the receptor tyrosine kinases (RTKs), using the pulse-chase endocytic recycling assay^51^. The cells transiently expressing Halo-tagged β2AR or EGFR (tagged at the extracellular N-terminus) were labeled using the cell-impermeable fluorogenic HaloTag ligand (JF_635_i-HaloTag ligand)^51^ and subsequently stimulated with their corresponding ligands, isoproterenol (ISO) or EGF, to stimulate dye-labeled Halo-β2AR or Halo-EGFR endocytosis. This allowed us to track the ligand-activated endocytosis process and subsequent intracellular trafficking and recycling of the fluorescently labeled receptors by TIRF microscopy. One to two minutes after β2AR or EGFR endocytosis, they became concentrated in vesicular or tubular structures that moved beneath the plasma membrane. Certain carriers containing Halo-β2AR or Halo-EGFR paused their movement, followed by a burst of VAMP2-pHluorin fluorescence, loss of receptor signal, and subsequent recruitment of the sensor (Fig. 8b,c). Approximately 40% of Halo-β2AR or Halo-EGFR exocytosis events recruited the PH(PLCδ1)-Aux1 sensor (Fig. 8b,c). Meanwhile, the fraction of sensor-labeled fusion events that contained the recycled β2AR and EGFR increased from the early to late stages of ligand stimulation (Fig. 8d, Extended Data Fig. 10b, and Supplementary Video 16). We also observed the recruitment of the sensor to the β2AR-enriched tubulovesicular carriers generated from EEA1-positive early endosomes (Extended Data Fig. 10c). These results collectively demonstrate the involvement of early endosome-derived clathrin-associated carriers in both the early and late stages of signaling receptor recycling.

An unexpected observation was that the activation of β2AR with ISO resulted in an increased frequency of fusion of clathrin-associated recycling carriers with the plasma membrane, which could be prevented by preincubating the cells with a β2AR antagonist (Fig. 8e). Similarly, EGF stimulation caused a transient enhancement in the fusion frequency in a concentration-dependent manner (Fig. 8f), and this enhancement was suppressed by EGFR depletion (Extended Data Fig. 10d). Inhibiting Arf1 activity effectively blocked the ISO- or EGF-induced increase in fusion event frequency (Fig. 8g,h). Consistent with these observations, we found that the removal of serum (and thus growth factors) from the culture medium reduced the fusion frequency of the carriers with the plasma membrane, whereas the addition of serum increased the frequency (Extended Data Fig. 10e). Collectively, these results indicate that the formation and dynamics of CARP can be modulated by cargo receptors and receptor-activated signaling.

To assess the functional relevance of clathrin-associated recycling carriers in regulating β2AR and TfR recycling, we interfered with CARP by inhibiting Arf1 activity through BFA or knocking down Rab1a/Rab1b expression by siRNA. These treatments impaired the recycling of β2AR and TfR (Extended Data Fig. 10f-i). To further evaluate the global effect of CARP depletion on cell surface proteomes, we analyzed the biotinylated cell surface proteins in cells treated with control siRNA or siRNA targeting Rab1a/Rab1b or Arf1 by mass spectrometry. We found that a wide array of integral membrane proteins was reduced from the surfaces of cells with Rab1a/Rab1b or Arf1 depletion (Fig. 8i). These proteins include receptors such as TfR (TFRC), IGF1R (insulin-like growth factor 1 receptor), and TLR2 (Toll-like receptor 2), and TMEM106B (an alternative receptor mediating SARS-CoV-2 infection)^52^. The identification of TfR validated the experimental approach. Several nutrient transporters, including SLC9A1 (sodium/hydrogen exchanger), SLC1A5 (neutral amino acid transporter), and SLC30A1 (zinc transporter), were also reduced from the cell surfaces after Rab1a/Rab1b and Arf1 depletion.

## Discussion

In this study, we have discovered and systematically characterized CARP, an endosomal recycling mechanism that involves the fast recycling of internalized receptors from early endosomes to the plasma membrane via a kiss-and-run fusion process (Fig. 8j). Clathrin, along with various regulatory proteins including AP1, Arf1, Rab1, and Rab11, is enriched at subregions on early endosomes lacking known retrieval complexes (Fig. 7h-p and Extended Data Fig. 9h-k). The resulting clathrin-associated tubulovesicular carriers are then transported to the plasma membrane with the assistance of actin (Fig. 7a,b and Extended Data Fig. 9a). Subsequently, these clathrin-associated carriers partially fuse with the plasma membrane (Fig. 2 and Extended Data Fig. 2), release endocytosed receptors (Fig. 8a-c and Extended Data Fig. 10a,b), pinch off (Fig. 6 and Extended Data Fig. 8), and then reenter the cells (Fig. 1h-k). Therefore, in contrast to the canonical Rab4-mediated fast recycling and Rab11-mediated slow recycling pathways, CARP is characterized by the employment of a distinct set of molecular machinery for generating the recycling carrier and facilitating the unexpected kiss-and-run membrane fusion (Fig. 8j).

The process of endosomal sorting requires the spatial enrichment of cargoes within existing or induced tubulovesicular subdomains on sorting endosomes, followed by membrane scission to generate transport carriers containing the cargoes destined for various membrane destinations^9^. Several distinct adaptor proteins, including AP1/AP3, GGAs, ACAP1, and Akt, are capable of recruiting clathrin to the endosomal membrane^53–55^. However, direct observation of vesicular carriers budding from clathrin-associated subdomains on early endosomes and their transportation directly to the plasma membrane has been lacking. The involvement of clathrin-associated subdomains on early endosomes in endocytic recycling remains enigmatic. By directly imaging the trafficking of clathrin-associated carriers from within cells to the plasma membrane, our results suggest that clathrin and adaptors can regulate the generation and transportation of recycling carriers independently of the canonical multimeric retrieval complexes. Previous studies have found that clathrin knockdown impaired the regeneration of synaptic vesicles at endosomes during ultrafast endocytosis at hippocampal synapses^56^. These observations suggest that, beyond its well-established role in endocytic vesicle formation, clathrin is likely involved in regulating the generation of recycling carriers at endosomes. On the other hand, given the canonical roles of Arf1 and AP1 in regulating trafficking between endosomes and the TGN^37^, it remains to be investigated whether TGN-derived carriers contribute to the generation of clathrin-associated recycling carriers from early endosomes (Fig. 8j).

Kiss-and-run is a mode of membrane fusion wherein the vesicle releases its contents through a transient fusion pore and is then retrieved at the site of fusion without fully collapsing into the plasma membrane^57,58^. Although still highly controversial, it has been suggested that synaptic vesicles can transiently fuse with the plasma membrane and release their contents by forming a narrow fusion pore, a process that occurs quickly (less than 1–2 seconds) and does not rely on clathrin^57, 58^. The kiss-and-run membrane fusion has also been observed during the exocytosis of post-Golgi vesicles and extensively studied during the release of secretory granules in non-neuronal secretory cells^29, 57, 58^. During the kiss-and-run exocytosis of post-Golgi vesicles, clathrin was found to associate with approximately half of the secretory vesicles to prevent their complete fusion^29^. However, whether endocytic recycling vesicles utilize full collapse, kiss-and-run, dynamic pore, or other mechanisms for efficient membrane fusion and cargo release has remained unknown. We found that clathrin was associated with a fraction of VAMP2-positive exocytosis events. These clathrin-associated recycling carriers exhibit kiss-and-run partial fusion at the plasma membrane, rather than instant complete collapse. In addition to clathrin, we observed that several other proteins, including the EHD proteins and clathrin-adaptor proteins CALM and EpsinR, were recruited to the partially fused recycling carriers at the plasma membrane. The dynamin-like EHD ATPases can form ring-like oligomers to induce/stabilize membrane bending and promote the tubulation and fission of lipid tubules^59^. Thus, clathrin, EHD proteins, or other proteins may form a barrier or stabilize the vesicles to prevent their full collapse. However, the precise mechanisms underlying kiss-and-run fusion of recycling vesicles, the mechanisms preventing clathrin from dissociating from the fused carriers, and how clathrin is organized on these carriers, require further investigation. Recent observations also demonstrate the kiss-and-run mechanism in cargo sorting from endocytic Rab5-positive vesicles to sorting endosomes and from tubular sorting endosomes to the Rab11-positive recycling vesicles^60, 61^. Thus, kiss-and-run could potentially serve as a general mechanism in the sorting and trafficking of endocytic recycling, enhancing the efficiency and robustness of the endocytic recycling system^60, 61^.

Given that ∼70–80% of internalized integral cell surface proteins are recycled back to the plasma membrane to maintain cellular functions and homeostasis^2, 3^, it is not surprising that multiple recycling pathways exist within cells. Our findings indicate that CARP is involved in the constitutive recycling process. Another intriguing finding was the ability of CARP to mediate the rapid recycling of signaling receptors back to the plasma membrane within three minutes after ligand-induced endocytosis. Currently, it remains unclear whether the recycling carriers have cargo specificity. Notably, the recycling carriers we identified appeared to lack sorting nexin proteins. The signaling receptors we examined were internalized through clathrin-mediated endocytosis, suggesting that binding to clathrin or its adaptors may be involved in cargo sorting and enrichment on early endosomes. This is supported by the observation that cargoes internalized via clathrin-dependent endocytosis were particularly enriched in Rab11-labeled recycling structures and segregated from cargoes internalized through clathrin-independent endocytosis^62^. Additionally, previous studies have reported that signaling receptors, such as β2AR and EGFR, can regulate the dynamics of clathrin-coated pits^63, 64^. Our findings suggest that recycling receptors can also affect the dynamics of recycling carriers. Activation of EGFR or β2AR signaling resulted in an increased frequency of fusion events with the plasma membrane, which was suppressed by inhibiting Arf1 activity. These findings align with previous observations that the activation of EGFR or the chemokine receptor CXCR4 of GPCR promoted Arf1 activation^65, 66^. Collectively, these results indicate that CARP can facilitate the fast and efficient recycling of receptors, a process which can be further enhanced by receptor-mediated signaling activation.

Interference with CARP exerted a global effect on the cell surface proteome, implying the broad involvement of CARP in regulating receptor and transporter recycling, cell signaling, and nutrient supply. Notably, the aberrant expression or activation of Rab1 and Arf1 has been linked to various human diseases, including cancer, neurodegenerative diseases, and neurodevelopmental disorders^67, 68^. Therefore, the discovery of CARP presents new opportunities for further exploring the physiological and pathological functions, as well as the regulatory mechanisms, of endocytic recycling.

## Methods

### Cell culture and chemical treatment

SUM159 cells^69^ were cultured at 37°C and 5% CO_2_ in DMEM/F12 (Corning or Gibco), supplemented with 5% FBS (Gibco), 100 U/ml penicillin-streptomycin (Corning), 1 μg/ml hydrocortisone (Sigma-Aldrich), 5 μg/ml insulin (Sigma-Aldrich), and 10 mM HEPES (Corning) at pH 7.4. COS7 cells were cultured at 37°C and 5% CO_2_ in DMEM (Corning) supplemented with 10% FBS, and 100 U/ml penicillin-streptomycin. U2OS cells were cultured at 37°C and 5% CO_2_ in McCoy’s 5A medium (Gibco) supplemented with 10% FBS and 100 U/ml penicillin-streptomycin. The cells were routinely verified to be mycoplasma-free using the TransDetect PCR Mycoplasma Detection Kit (TransGen Biotech).

To starve cells, cells were washed three times with DPBS (Corning) and then cultured in DMEM/F12 (Corning) for the indicated times. Actin and microtubule networks were disrupted by treating cells with latrunculin A (Cayman; 0.5 μM for 30-60 min), blebbistatin (Targetmol; 25 μM for 2-3 h), or nocodazole (Sigma-Aldrich; 10 μM for 1-2 h) in the SUM159 culture medium. To inhibit Arf activity, cells were treated with BFA (MedChemExpress; 10 μg/ml for 30 min) or EXO2^70^ (Sigma-Aldrich; 50 μM for 10-30 min) in the SUM159 culture medium. To inhibit dynamin activity, cells were treated with Dynole 2-24^71^ (Sigma- Aldrich) in serum-free DMEM/F12 (5 μM for 5-20 min). Rapamycin (Merck Millipore; 0.5 μM) was added to cells during continuous imaging using a syringe pump (Harvard Apparatus). JF_646_-HaloTag ligand was purchased from Promega. JFX_650_-HaloTag ligand, JF_635_i-HaloTag ligand, and JF_549_i-SNAP-tag ligand were kind gifts from Dr. Luke D. Lavis (Janelia Research Campus)^51, 72^.

### Plasmids and transfection

The DNA sequences encoding various proteins were amplified by PCR from cDNA clones or cDNA of SUM159/HEK293 cells, and then inserted into a vector containing mScarlet-I to generate the plasmids used for imaging-based screening. A (GGS)_5_ linker (5’-GGAGGATCCGGTGGATCTGGAGGTTCTGGTGGTTCTGGTGGTTCC-3’) was placed between mScarlet-I and the amplified DNA sequence. The EGFP-Tubby_c_-Aux1 sensor was constructed by substituting PH(PLCδ1) for Tubby_c_ in EGFP-PH(PLCδ1)-Aux1. Str-KDEL-TfR1-SBP-mCherry was constructed based on the construct Str-KDEL-SBP-mCherry-CD-MPR^73^. The plasmids generated in this study and those obtained from Addgene are listed in Supplementary Table 1. Cells were transfected using Lipofectamine 3000 (Invitrogen) according to the manufacturer’s instructions.

### siRNA knockdown

Cells were transfected with gene-specific siRNA oligos or a mixture of non-targeting control oligos using Lipofectamine RNAiMAX (Invitrogen) following the manufacturer’s instructions. Knockdown of the expression of dynamin2, Rab1a, or Rab1b was achieved by transfecting the cells with gene-specific siRNA oligos for 72 h. Knockdown of other proteins was achieved by two sequential transfections, the first one on day one (after overnight plating) and the second one on day three, followed by imaging analysis on day five. The siRNA sequences used are listed in Supplementary Table 2. The knockdown efficiency was verified by western blot analysis or real-time quantitative PCR. Total RNA was extracted using Eastep^®^ Super Total RNA Extraction Kit (Promega). The PrimeScript RT Master Mix Kit (TaKaRa) was used to generate cDNA from the extracted RNA. Real-time quantitative PCR was performed using SYBR^®^ Green Realtime PCR Master Mix (Toyobo) and the CFX Connect Real-Time PCR Detection System (Bio-Rad). GAPDH expression was analyzed for normalization. The primers used for real-time quantitative PCR are listed in Supplementary Table 3.

### Generation of knock-in cell lines

SUM159 cells were gene-edited to incorporate mScarlet-I, miRFP670nano, or HaloTag into the C-terminus of AP2σ2, CLTA, or dynamin2 using a TALEN-based protocol as described^11, 74, 75^. SUM159 cells were gene-edited to incorporate mEGFP, mScarlet-I, mScarlet3, TagRFP, or HaloTag into the N- or C-terminus of other targeted genes using the CRISPR/Cas9 approach as described^11, 76^. The gRNA sequences utilized for generating the knock-in cell lines are listed in Supplementary Table 4. The sgRNA sequences were cloned into the pSpCas9(BB)-2A-Puro (PX459) plasmid (Addgene).

Donor constructs for homologous recombination were generated by cloning into the pUC19 vector with two ∼600-800-nucleotide fragments of genomic DNA upstream and downstream of the start or stop codon of the targeted gene, along with the open reading frame of mEGFP/mScarlet-I/mScarlet3/TagRFP/HaloTag, using the pEASY-Uni Seamless Cloning and Assembly Kit (TransGen Biotech). A (GGS)_3_ linker (5’- GGAGGTTCTGGTGGTTCTGGTGGTTCC-3’) was inserted between the start or stop codon of the targeted gene and the open reading frame of mEGFP/mScarlet-I/mScarlet3/TagRFP/HaloTag.

SUM159 cells were transfected with the donor plasmid and the PX459 plasmid containing the sgRNA targeting sequence using Lipofectamine 3000 (Invitrogen). Five to seven days after transfection, cells expressing mEGFP, mScarlet-I, mScarlet3, TagRFP, or HaloTag (Promega) were enriched by fluorescence-activated cell sorting (FACS) (FACSAria II, BD Biosciences). The HaloTag-expressing cells were stained with 20 nM of JF_549_-HaloTag ligand for 10 min in the culture medium, followed by three washes with DPBS before sorting. The sorted cells were expanded and subsequently subjected to either single-cell sorting into 96-well plates or bulk sorting to get a pool of positive cells. The monoclonal cells were screened for successful incorporation in the genomic locus of the fluorescent tag by PCR using GoTaq DNA Polymerase (Promega). The genome-edited cells were confirmed through imaging, PCR, and western blot analysis.

### Generation of knockout cell lines

Knockout of targeted genes in SUM159 cells was performed using the CRISPR/Cas9 approach as described^76^. The sgRNA sequence targeting the desired gene was cloned into either pSpCas9(BB)-2A-GFP (PX458) (Addgene) or pSpCas9(BB)-2A-Halo. The sgRNA sequences used for knockout of Rab11b^77^, VPS35^78^, SNX27^78^, VPS29^79^, CCDC22^79^, SNX17^80^, SNX1^81^, SNX2^81^, SNX5^81^, SNX6^81^, AP1μ1^82^, Arf1, or AP2σ2 are listed in Supplementary Table 5. SUM159 cells were transfected with the constructed plasmid using Lipofectamine 3000. Cells expressing GFP or Halo (labeled by JF_549_-HaloTag ligand) were subjected to single-cell sorting into 96-well plates 24-36 h after transfection (FACSAria II, BD Biosciences). Monoclonal cell populations with mutations in both alleles were identified through sequencing, and the loss of targeted protein expression was confirmed by western blot analysis.

### Generation of stable cell lines

SUM159, COS7, and U2OS cells stably expressing EGFP-PH(PLCδ1)-Aux1, VAMP2-pHluorin, or α- TagRFP-AP2 were generated by transduction with the pHAGE lentiviral vector encoding EGFP- PH(PLCδ1)-Aux1, or VAMP2-pHluorin, or α-TagRFP-AP2. The α-TagRFP-AP2 construct was generated based on the α-EGFP-AP2 construct described in the previous study^18^. Pools of positive cells were collected by FACS 5-7 days after transduction. COS7 or U2OS cells stably expressing relatively low levels of EGFP- PH(PLCδ1)-Aux1 were enriched by one additional bulk sorting. Clonal SUM159 cells expressing relatively low levels of the sensor were obtained through single-cell sorting.

### Live-cell imaging using TIRF microscopy

The bottom surfaces of cells were imaged using a TIRF microscope built either on a Nikon Ti2-E microscope or a Nikon TiE microscope. The Nikon Ti2-E microscope was equipped with a manual TIRF Illuminator Unit (Nikon), a CFI Apochromat TIRF 100X objective (1.49 NA, Nikon), a Perfect Focus Unit (Nikon), a UNO Stage Top Incubator (Okolab), OBIS CellX lasers (405, 488, 561 and 637 nm, Coherent), W-VIEW GEMINI-2C Image Splitting Optics (Hamamatsu), and two EMCCD cameras (Evolve 512 Delta, Photometrics). The integrated 2× tube lens on the Nikon Ti2-E microscope was applied to achieve the final pixel size corresponding to 80 nm of the image. Images were acquired using Micro-Manager 1.4^83^. The Nikon TiE microscope was equipped with a motorized TIRF Illuminator Unit (Nikon), a CFI Apochromat TIRF 100X objective (1.49 NA, Nikon), a Perfect Focus Unit (Nikon), a Motorized XY stage (Prior Scientific), a fully enclosed and environmentally controlled cage incubator (Okolab), OBIS 488, 561 and 647 nm lasers (Coherent), W-VIEW GEMINI Image splitting optics (Hamamatsu), and an EMCCD camera (iXon Life 888, Andor Technology). The 1.5× tube lens integrated on the Nikon TiE microscope was applied to achieve the final pixel size corresponding to 86.7 nm of the image. Imaging sequences were acquired using Micro-Manager 2.0^83^.

The cells were plated on single-well, 4-well, or 8-well confocal dishes (Cellvis) approximately 6-8 hours after transfection. Cells expressing relatively low levels of the sensor or other proteins were imaged at the bottom surfaces by TIRF microscopy every 1 or 1.5 s for a duration of 3 or 5 min in phenol-free DMEM/F12 (Corning) containing 5% FBS and 20 mM HEPES. The variable-angle TIRF was performed using the Nikon TiE microscope with a motorized TIRF Illuminator Unit. Montages were created in Fiji^84^. Kymographs were created using the Multi Kymograph plugin in Fiji.

### Identification of fusion events at the plasma membrane of cells imaged by TIRF microscopy

The identification of AP2-negative clathrin-associated recycling events that recruited the PH(PLCδ1)-Aux1 sensor in each time series was done in the following steps. The cmeAnalysis software package^17^ (https://github.com/DanuserLab/cmeAnalysis) was used to detect and track the EGFP-PH(PLCδ1)-Aux1 sensor along with the associated clathrin and AP2, with the sensor channel as the ’master’ channel and the AP2/clathrin channels as the ’slave’ channels. The standard deviation of the estimated Gaussian point spread function was 1.43 pixels for EGFP, 1.58 pixels for TagRFP, and 1.61 pixels for 670nano. The minimum and maximum tracking search radii were 1 and 3 pixels, and the maximum gap length in a trajectory was 1 frame. Valid trajectories with lifetimes longer than 5 s, less than 20% of gap frames, a mean signal-to-background ratio above 1.08, maximum intensity 1.3 times higher than the intensity of the first frame, and statistically significant clathrin signal but undetectable AP2 signal in the slave channels, were automatically selected. Intensity-lifetime cohort plots were generated as described (Aguet et al., 2013). The fusion event frequency (events/min/μm^2^) for each cell was determined by dividing the total number of fusion events detected in the time series by the movie length (3 or 5 minutes) and cell area. The number of cells used for calculating the fusion frequency is reported in figure legends.

To generate the membrane fusion frequency heatmap of clathrin-associated recycling carriers, each cell was first normalized into a circular shape based on the centroid position and cell boundary. Then, the relative position of each fusion event was recalculated after normalization. The positions of fusion events from multiple cells were combined to generate the fusion frequency heatmap in MATLAB (MathWorks).

The detection and tracking of VAMP2-pHluorin fusion events were performed using the cmeAnalysis software package^17^. The minimum and maximum tracking search radii were set to 1 and 3 pixels, and the maximum gap length in a trajectory was 1 frame. For valid tracks with less than 20% of gap frames and maximum fluorescence intensity (F1) 1.4 times higher than the mean intensity of all frames, the difference between F1 and the average intensity from three early-stage frames starting three frames before the peak (F0) was calculated (F1-F0). Tracks with F1-F0 greater than F0 by a ratio of 0.3 were selected automatically and verified manually. Tracks with peak frames appearing in the first five frames were also selected automatically and verified manually. Tracks with significant signals from the slave channel were used for further analysis.

The identification of Rab11a/Rab11b spots in each frame of the time series was performed by TrackMate 7 (Fiji)^85, 86^ (https://github.com/fiji/TrackMate). The density of Rab11a/Rab11b spots (spots/frames/μm^2^) for each cell was determined by dividing the total number of Rab11a/Rab11b spots detected in the time series by the movie length and cell area.

### Live-cell imaging using spinning-disk confocal microscopy

The spinning-disk confocal microscope was built on the Nikon TiE microscope described above. The TiE microscope was also equipped with a CSU-X1 spinning-disk confocal unit (Yokogawa) and an EMCCD camera (iXon Ultra 897, Andor Technology) or an sCMOS camera (Aries 16, Tucsen Photonics) mounted on the left side port. To monitor the dynamic movement of intracellular clathrin-associated carriers to the plasma membrane, cells were plated in single-well confocal dishes (Cellvis) at high density. The following day, cells reached confluence to minimize dynamic lateral movement of the dorsal surface during imaging. Cells were imaged every 1.5 s at the middle plane to facilitate visualization and tracking of the dynamic movement of intracellular clathrin-associated carriers toward the cell membrane between the cell-cell contacts. The plasma membrane was segmented manually in Fiji. For time-lapse imaging at the bottom surface, individual cells were imaged every 1.5 or 2 s for a duration of 3 or 5 min in phenol-free DMEM/F12 (Corning) containing 5% FBS and 20 mM HEPES. The clathrin-associated carriers or EEA1-labeled endosomes were detected and tracked by TrackMate 7.

### Live-cell imaging with SIM

Live-cell SIM imaging was performed on the Multi-SIM system as described^87^. Cells were imaged in a fully enclosed and environmentally controlled cage incubator (Okolab) with a CFI SR HP Apo TIRF 100X objective (1.49 NA, Nikon), and an sCMOS camera (Kinetix, Photometrics). Live-cell imaging of clathrin-associated carriers in CLTA-TagRFP^+/+^ AP2-Halo^+/+^ cells stably expressing EGFP-sensor was conducted using the grazing incidence SIM (GI-SIM). The clathrin channel of the SIM images underwent additional processing using the rationalized deep learning algorithm to reduce the noise/background-induced reconstruction artifacts^87^. To measure the relative distribution of Arf1 with respect to other proteins on individual EEA1-positive early endosomes, the cells were imaged using the low-NA GI-SIM and results were shown by polar plots. These plots were generated based on the EEA1-positive early endosomes containing both Arf1 and other proteins. The distances between each protein and the center of the endosomes, as well as the angles between Arf1 and each protein, were measured by Fiji. The polar plots were then generated using the polarscatter function in MATLAB.

### Correlative fluorescence and FIB-SEM imaging

For correlative fluorescence and FIB-SEM imaging, cells were plated on 35 mm glass-bottom dishes with position markers (P35G-1.5-14-C-GRID, MatTek) for 24 h and then fixed with 4% PFA. The cells were first imaged through spinning-disk confocal microscopy. The dish position markers were recorded using bright-field imaging. Immediately after imaging, the cells were fixed with 2.5% glutaraldehyde for 1 hour at room temperature. The samples were post-fixed by 2% osmium tetroxide/1.5% potassium ferrocyanide, followed by 1% thiocarbohydrazide and 1% osmium. The samples were then stained with 1% aqueous uranyl acetate, dehydrated through a series of graded ethyl alcohol (50%, 70%, 80%, 90%, 95%, and 100%), and embedded in SPI-Pon 812 resin (SPI Supplies). The cell samples were then imaged volumetrically in 3D using an electron microscope that combines a focused ion beam (FIB) and a high-resolution field emission scanning electron microscope (SEM) (Helios Nanolab G3 UC, Thermo Fisher Scientific). The milling and imaging processes were sequentially repeated and long series of images (600) were acquired through a fully automated procedure. The image resolution on the XY plane was approximately 3 nm/pixel. The resolution on the Z axis was 5 nm, which corresponds to the thickness of the material layer removed by FIB. The final image was captured at a pixel size of 6144×4096, providing a field of view of approximately 18×12 μm, equivalent to a magnification of 15000. Amira software (Thermo Fisher Scientific) was used for automatic alignment, image denoising, and signal normalization across slices. Based on the position markers (letters) on the resin block and those recorded in bright-field imaging, the corresponding fluorescence image of each cell was identified. Initially, the position, rotation angle, and scaling were determined using the fluorescence image and FIB-SEM image of the nucleus-containing middle plane of the cell in Fiji. Further adjustments were made by aligning the positions of AP2-positive clathrin-coated pits at the plasma membrane in the fluorescence image with those structures in the FIB-SEM image. After refining the alignment of the fluorescence and FIB-SEM images, the structures in the FIB-SEM image corresponding to the AP2-negative sensor-positive clathrin-containing structures identified in fluorescence images were segmented and reconstructed in Amira.

### Live-cell imaging with lattice light-sheet microscopy

CLTA-Halo^+/+^ AP2-TagRFP^+/+^ cells stably expressing EGFP-sensor were imaged using the ZEISS Lattice Lightsheet 7 microscope. The Sinc3 15 x 650 lattice light sheet was used for imaging. The 3D volume of the cell was recorded by scanning the cell (40 slices) every ∼3.9 s for 3 min, with a step size of 145 nm along the Z-axis. Fluorescence signals were collected using a detection objective (44.83x / NA 1.0) and an sCMOS camera (pco.edge 4.2). The cells were imaged at 37°C and 5% CO_2_ in phenol-free DMEM/F12 (Corning) containing 5% FBS and 20 mM HEPES. The collected raw images were reconstructed and deconvoluted using ZEN 3.8 (ZEISS).

### Measurement of receptor recycling

TfR recycling under BFA treatment or Rab1a/Rab1b knockdown was measured by a modified imaging-based assay as described^88^. Briefly, SUM159 cells were serum-starved for 2 h in DMEM/F12 containing 20 mM HEPES and then incubated on ice for 10 min with 5 μg/mL Alexa Fluor 568-conjugated transferrin (Thermo Fisher Scientific). After the incubation, the cells were washed with ice-chilled PBS and then transferred to the pre-warmed culture medium at 37°C for the indicated times. Cells were then briefly washed (2 min) with ice-chilled acid wash medium (150 mM NaCl, 1 mM MgCl_2_, 0.125 mM CaCl_2_, and 0.1 M glycine, pH 2.5) for two times and fixed with 4% paraformaldehyde for 20 min at room temperature.

β2AR recycling under BFA treatment or Rab1a/Rab1b knockdown was evaluated by a modified imaging-based assay as described^89^. Briefly, SUM159 cells stably expressing SNAP-β2AR (SNAP-tag at the extracellular N-terminus) were incubated with the cell-impermeable JF_549_i-SNAP-tag ligand in the culture medium at 37°C for 10 min. After washing with PBS, the cells were treated with ISO (10 μM for 10 min) to promote the internalization of dye-labeled receptors. The cells were then fixed or further incubated with the medium containing the β-blocker propranolol for an additional 20 min before fixation. The cells were imaged in PBS using spinning-disk confocal microscopy.

The live-cell tracking of the endocytic recycling of Halo-β2AR and Halo-EGFR was performed with pulse-chase assays using the JF_635_i-HaloTag ligand as described^51^. Briefly, cells stably expressing VAMP2- pHluorin and mRuby3-sensor were transiently transfected with Halo-β2AR or Halo-EGFR, and then stained with membrane-impermeable JF_635_i-HaloTag ligand in culture medium (40 nM for 10 min). After three washes with culture medium, the cells were treated with ISO (10 μM) or EGF (100 ng/ml) for 3 min to stimulate the endocytosis of dye-labeled Halo-β2AR or Halo-EGFR. Then the cells were imaged by TIRF microscopy to capture the exocytosis of internalized receptors.

### Cell surface proteomics

Cell surface protein biotinylation and isolation were performed using the Pierce™ Cell Surface Protein Biotinylation and Isolation Kit (Thermo Fisher Scientific, A44390) according to the manufacturer’s instructions. Briefly, cells were treated with control siRNA or siRNA targeting Rab1a/b or Arf1 for 72 h. The cells were washed with PBS and then incubated with Sulfo-NHS-SS-Biotin at room temperature for 10 min. After removing the labeling solution, the cells were washed with ice-cold TBS twice, scarped into ice-cold TBS, and then centrifuged at 500 *g* for 3 min at 37°C. The cell pellet was resuspended in the lysis buffer with protease inhibitors and pipetted up and down 20 times, incubated on ice for 30 min with a brief vertexing at the beginning and end of the incubation, and then centrifuged at 15,000 *g* for 5 min at 4°C. The supernatant was added to the NeutrAvidin™ Agarose pretreated column and then incubated for 30 min at room temperature on a rotator. After four times washing with the Wash Buffer, the column was incubated with the prepared Elution buffer for 30 min at room temperature on a rotator. Proteins were collected in a collection tube after centrifuging the column for 2 min at 1,000 g. Proteins were separated on SDS-PAGE for subsequent in-gel digestion. LC-MS analyses of peptide samples were performed on a hybrid ion trap-Orbitrap mass spectrometer (LTQ-Orbitrap Velos, Thermo Scientific) coupled with nanoflow reversed-phase liquid chromatography (EASY-nLC 1200, Thermo Scientific). The raw MS files were searched against the UniProt human protein database (Proteome IP UP000005640, version 2021_04) using Mascot software (v2.3.02, Matrix Science). Results from three independent experiments are shown in Supplementary Table 6. Membrane proteins with averaged fold changes > 1.2 from three independent experiments were considered as candidates and shown in the figure.

### Western blot analysis

The cells were solubilized in RIPA lysis buffer (Sigma-Aldrich) with a protease inhibitor cocktail (Pierce) for 10 min on ice and then pelleted at 13,400 g for 15 min at 4 °C. The supernatant was mixed with 5× sample buffer (GenScript), heated to 100°C for 10 min, and then fractionated by SDS–PAGE (Vazyme) and transferred to nitrocellulose membranes (Cell Signaling). The membranes were incubated in TBST buffer containing 5% skim milk for 1 h at room temperature, followed by overnight incubation at 4°C with the specific primary antibodies. Following three washes with TBST (5 min each), the membranes were incubated with the appropriate HRP-conjugated secondary antibody (Beyotime, 1:1,000) for 1 h at room temperature. The membranes were then incubated with the SignalFire^TM^ ECL Reagent (Cell Signaling) and imaged by the Tanon-5200 Chemiluminescent Imaging System (Tanon) or MiniChemi 610 Chemiluminescent Imaging System (SINSAGE). The primary antibodies used in this study are listed in Supplementary Table 7.

### Statistical tests

Statistical analyses were performed using GraphPad Prism 9 (GraphPad Software). The unpaired, two-tailed Student’s *t*-test was used to compare the two groups. The ordinary one-way ANOVA, followed by Tukey’s multiple-comparisons test, was used to compare three or four groups. *P* < 0.05 was considered statistically significant. No statistical methods were used to predetermine sample sizes.

### Data availability

The data that support the findings of the current study are included in Extended Data Figs. 1–10, Extended Tables 1-7, and Supplementary Videos 1–16. The source imaging data presented in this paper are available from the corresponding author upon request. The full gel source data for western blot and PCR have been provided in Supplementary Figs. 1 and 2.

### Code availability

The download links for published codes were provided in the Methods. Other custom MATLAB codes are available upon request from the corresponding author.

## Acknowledgments

We thank Dr. Tom Kirchhausen for critical reading and discussions, and TALEN constructs used for genome-editing of AP2S1, CLTA, and dynamin2. We thank Dr. Luke D. Lavis for the generous gifts of the HaloTag and SNAP-tag ligands. We thank Dr. Joan Brugge for generously providing SUM159 cells. We express our gratitude to Dr. Baoliang Song, Dr. Juan S. Bonifacino, and Dr. Zhiming Chen for generously providing the constructs EGFP-myosin Vb, Str-KDEL-SBP-mCherry-CD-MPR, and the internally EGFP-tagged α-adaptin, respectively. We thank Dr. Yu Luo, Wenyi Shen, Wenhao Dong, and Guangyuan Zhong from ZEISS China and ZEISS Microscopy Customer Center Beijing for their technical support in collecting and analyzing imaging data on Lattice Lightsheet 7. We thank Dr. Isabel Hanson for editing. This work was supported by grants from the National Natural Science Foundation of China (92354305 and 32321004 to K.H; 92154001 to A.S), the Ministry of Science and Technology of the People’s Republic of China (2022YFA1304500 and 2021YFA0804802 to K.H.), the Youth Innovation Promotion Association CAS (2023105 to N.L.), and the State Key Laboratory of Molecular Developmental Biology of China.

## Competing interests

The authors declare no competing interests.

## Notes

### Competing Interest Statement

The authors have declared no competing interest.

## References

1. Schmid, S.L., Sorkin, A. & Zerial, M. Endocytosis: Past, present, and future. Cold Spring Harb Perspect Biol 6, a022509 (2014).

2. Solinger, J.A. & Spang, A. Sorting of cargo in the tubular endosomal network. Bioessays 44, e2200158 (2022).

3. Cullen, P.J. & Steinberg, F. To degrade or not to degrade: mechanisms and significance of endocytic recycling. Nat Rev Mol Cell Biol 19, 679–696 (2018).

4. Mellman, I. & Yarden, Y. Endocytosis and cancer. Cold Spring Harb Perspect Biol 5, a016949 (2013).

5. Schreij, A.M., Fon, E.A. & McPherson, P.S. Endocytic membrane trafficking and neurodegenerative disease. Cell Mol Life Sci 73, 1529–1545 (2016).

6. Klumperman, J. & Raposo, G. The complex ultrastructure of the endolysosomal system. Cold Spring Harb Perspect Biol 6, a016857 (2014).

7. Jovic, M., Sharma, M., Rahajeng, J. & Caplan, S. The early endosome: a busy sorting station for proteins at the crossroads. Histol Histopathol 25, 99–112 (2010).

8. Grant, B.D. & Donaldson, J.G. Pathways and mechanisms of endocytic recycling. Nature reviews. Molecular cell biology 10, 597–608 (2009).

9. McNally, K.E. & Cullen, P.J. Endosomal Retrieval of Cargo: Retromer Is Not Alone. Trends Cell Biol 28, 807–822 (2018).

10. Hsu, V.W., Bai, M. & Li, J. Getting active: protein sorting in endocytic recycling. Nat Rev Mol Cell Biol 13, 323–328 (2012).

11. He, K. et al. Dynamics of phosphoinositide conversion in clathrin-mediated endocytic traffic. Nature 552, 410–414 (2017).

12. Chen, Z. & Schmid, S.L. Evolving models for assembling and shaping clathrin-coated pits. J Cell Biol 219 (2020).

13. Kirchhausen, T., Owen, D. & Harrison, S.C. Molecular structure, function, and dynamics of clathrin-mediated membrane traffic. Cold Spring Harb Perspect Biol 6, a016725 (2014).

14. Balla, T. Phosphoinositides: tiny lipids with giant impact on cell regulation. Physiol Rev 93, 1019–1137 (2013).

15. Pacheco, J., Cassidy, A.C., Zewe, J.P., Wills, R.C. & Hammond, G.R.V. PI(4,5)P2 diffuses freely in the plasma membrane even within high-density effector protein complexes. J Cell Biol 222 (2023).

16. Hammond, G.R.V., Ricci, M.M.C., Weckerly, C.C. & Wills, R.C. An update on genetically encoded lipid biosensors. Mol Biol Cell 33 (2022).

17. Aguet, F., Antonescu, C.N., Mettlen, M., Schmid, S.L. & Danuser, G. Advances in analysis of low signal-to-noise images link dynamin and AP2 to the functions of an endocytic checkpoint. Dev Cell 26, 279–291 (2013).

18. Mino, R.E., Chen, Z., Mettlen, M. & Schmid, S.L. An internally eGFP-tagged alpha-adaptin is a fully functional and improved fiduciary marker for clathrin-coated pit dynamics. Traffic 21, 603–616 (2020).

19. Lanni, F., Waggoner, A.S. & Taylor, D.L. Structural organization of interphase 3T3 fibroblasts studied by total internal reflection fluorescence microscopy. J Cell Biol 100, 1091–1102 (1985).

20. Kural, C. et al. Dynamics of intracellular clathrin/AP1- and clathrin/AP3-containing carriers. Cell Rep 2, 1111–1119 (2012).

21. Dubuke, M.L. & Munson, M. The Secret Life of Tethers: The Role of Tethering Factors in SNARE Complex Regulation. Front Cell Dev Biol 4, 42 (2016).

22. Ahmed, S.M. et al. Exocyst dynamics during vesicle tethering and fusion. Nature communications 9, 5140 (2018).

23. Mei, K. & Guo, W. The exocyst complex. Curr Biol 28, R922–R925 (2018).

24. Dingjan, I. et al. Endosomal and Phagosomal SNAREs. Physiol Rev 98, 1465–1492 (2018).

25. Chen, H., Weinberg, Z.Y., Kumar, G.A. & Puthenveedu, M.A. Vesicle-associated membrane protein 2 is a cargo-selective v-SNARE for a subset of GPCRs. J Cell Biol 222 (2023).

26. Miesenbock, G., De Angelis, D.A. & Rothman, J.E. Visualizing secretion and synaptic transmission with pH-sensitive green fluorescent proteins. Nature 394, 192–195 (1998).

27. Zhao, Y. & Keen, J.H. Gyrating clathrin: highly dynamic clathrin structures involved in rapid receptor recycling. Traffic 9, 2253–2264 (2008).

28. Luo, Y., Zhan, Y. & Keen, J.H. Arf6 regulation of Gyrating-clathrin. Traffic 14, 97–106 (2013).

29. Jaiswal, J.K., Rivera, V.M. & Simon, S.M. Exocytosis of Post-Golgi Vesicles Is Regulated by Components of the Endocytic Machinery. Cell 137, 1308–1319 (2009).

30. Pascolutti, R. et al. Molecularly Distinct Clathrin-Coated Pits Differentially Impact EGFR Fate and Signaling. Cell Rep 27, 3049–3061 e3046 (2019).

31. Presley, J.F. et al. ER-to-Golgi transport visualized in living cells. Nature 389, 81–85 (1997).

32. Boncompain, G. et al. Synchronization of secretory protein traffic in populations of cells. Nat Methods 9, 493–498 (2012).

33. Wandinger-Ness, A. & Zerial, M. Rab proteins and the compartmentalization of the endosomal system. Cold Spring Harb Perspect Biol 6, a022616 (2014).

34. Takahashi, S. et al. Rab11 regulates exocytosis of recycling vesicles at the plasma membrane. J Cell Sci 125, 4049–4057 (2012).

35. Monetta, P., Slavin, I., Romero, N. & Alvarez, C. Rab1b interacts with GBF1 and modulates both ARF1 dynamics and COPI association. Mol Biol Cell 18, 2400–2410 (2007).

36. Hatoyama, Y., Homma, Y., Hiragi, S. & Fukuda, M. Establishment and analysis of conditional Rab1- and Rab5-knockout cells using the auxin-inducible degron system. J Cell Sci 134 (2021).

37. Adarska, P., Wong-Dilworth, L. & Bottanelli, F. ARF GTPases and Their Ubiquitous Role in Intracellular Trafficking Beyond the Golgi. Front Cell Dev Biol 9, 679046 (2021).

38. Highland, C.M. & Fromme, J.C. Arf1 directly recruits the Pik1-Frq1 PI4K complex to regulate the final stages of Golgi maturation. Mol Biol Cell 32, 1064–1080 (2021).

39. Wang, J. et al. PI4P promotes the recruitment of the GGA adaptor proteins to the trans-Golgi network and regulates their recognition of the ubiquitin sorting signal. Mol Biol Cell 18, 2646–2655 (2007).

40. Henmi, Y. et al. PtdIns4KIIalpha generates endosomal PtdIns(4)P and is required for receptor sorting at early endosomes. Mol Biol Cell 27, 990–1001 (2016).

41. Mills, I.G. et al. EpsinR: an AP1/clathrin interacting protein involved in vesicle trafficking. J Cell Biol 160, 213–222 (2003).

42. Antonny, B. et al. Membrane fission by dynamin: what we know and what we need to know. EMBO J 35, 2270–2284 (2016).

43. Loerke, D. et al. Cargo and dynamin regulate clathrin-coated pit maturation. PLoS Biol 7, e57 (2009).

44. Mooren, O.L., Galletta, B.J. & Cooper, J.A. Roles for actin assembly in endocytosis. Annu Rev Biochem 81, 661–686 (2012).

45. Anitei, M. & Hoflack, B. Bridging membrane and cytoskeleton dynamics in the secretory and endocytic pathways. Nat Cell Biol 14, 11–19 (2011).

46. Chu, B.B. et al. Requirement of myosin Vb.Rab11a.Rab11-FIP2 complex in cholesterol-regulated translocation of NPC1L1 to the cell surface. J Biol Chem 284, 22481-22490 (2009).

47. Simonetti, B. & Cullen, P.J. Actin-dependent endosomal receptor recycling. Curr Opin Cell Biol 56, 22–33 (2019).

48. Kondo, Y. et al. ARF1 and ARF3 are required for the integrity of recycling endosomes and the recycling pathway. Cell Struct Funct 37, 141–154 (2012).

49. Wang, J. et al. Endosomal receptor trafficking: Retromer and beyond. Traffic 19, 578–590 (2018).

50. Nielsen, E., Severin, F., Backer, J.M., Hyman, A.A. & Zerial, M. Rab5 regulates motility of early endosomes on microtubules. Nature cell biology 1, 376–382 (1999).

51. Jonker, C.T.H. et al. Accurate measurement of fast endocytic recycling kinetics in real time. J Cell Sci 133 (2020).

52. Baggen, J. et al. TMEM106B is a receptor mediating ACE2-independent SARS-CoV-2 cell entry. Cell (2023).

53. D’Souza, R.S. et al. Rab4 orchestrates a small GTPase cascade for recruitment of adaptor proteins to early endosomes. Curr Biol 24, 1187–1198 (2014).

54. Hsu, J.W. et al. The protein kinase Akt acts as a coat adaptor in endocytic recycling. Nat Cell Biol 22, 927–933 (2020).

55. Li, J. et al. An ACAP1-containing clathrin coat complex for endocytic recycling. J Cell Biol 178, 453–464 (2007).

56. Watanabe, S. et al. Clathrin regenerates synaptic vesicles from endosomes. Nature 515, 228–233 (2014).

57. Alabi, A.A. & Tsien, R.W. Perspectives on kiss-and-run: role in exocytosis, endocytosis, and neurotransmission. Annu Rev Physiol 75, 393–422 (2013).

58. Chanaday, N.L., Cousin, M.A., Milosevic, I., Watanabe, S. & Morgan, J.R. The Synaptic Vesicle Cycle Revisited: New Insights into the Modes and Mechanisms. J Neurosci 39, 8209–8216 (2019).

59. Dhawan, K., Naslavsky, N. & Caplan, S. Sorting nexin 17 (SNX17) links endosomal sorting to Eps15 homology domain protein 1 (EHD1)-mediated fission machinery. J Biol Chem 295, 3837–3850 (2020).

60. Solinger, J.A., Rashid, H.O. & Spang, A. FERARI and cargo adaptors coordinate cargo flow through sorting endosomes. Nat Commun 13, 4620 (2022).

61. Solinger, J.A., Rashid, H.O., Prescianotto-Baschong, C. & Spang, A. FERARI is required for Rab11-dependent endocytic recycling. Nat Cell Biol 22, 213–224 (2020).

62. Xie, S. et al. The endocytic recycling compartment maintains cargo segregation acquired upon exit from the sorting endosome. Mol Biol Cell 27, 108–126 (2016).

63. Puthenveedu, M.A. & von Zastrow, M. Cargo regulates clathrin-coated pit dynamics. Cell 127, 113–124 (2006).

64. Alfonzo-Mendez, M.A., Sochacki, K.A., Strub, M.P. & Taraska, J.W. Dual clathrin and integrin signaling systems regulate growth factor receptor activation. Nat Commun 13, 905 (2022).

65. Khater, M., Bryant, C.N. & Wu, G. Gbetagamma translocation to the Golgi apparatus activates ARF1 to spatiotemporally regulate G protein-coupled receptor signaling to MAPK. J Biol Chem 296, 100805 (2021).

66. Boulay, P.L., Cotton, M., Melancon, P. & Claing, A. ADP-ribosylation factor 1 controls the activation of the phosphatidylinositol 3-kinase pathway to regulate epidermal growth factor- dependent growth and migration of breast cancer cells. J Biol Chem 283, 36425–36434 (2008).

67. Ishida, M. et al. A neurodevelopmental disorder associated with an activating de novo missense variant in ARF1. Hum Mol Genet 32, 1162–1174 (2023).

68. Yang, X.Z. et al. Rab1 in cell signaling, cancer and other diseases. Oncogene 35, 5699–5704 (2016).

## References

69. Forozan, F. et al. Molecular cytogenetic analysis of 11 new breast cancer cell lines. Br J Cancer 81, 1328–1334 (1999).

70. Feng, Y. et al. Retrograde transport of cholera toxin from the plasma membrane to the endoplasmic reticulum requires the trans-Golgi network but not the Golgi apparatus in Exo2-treated cells. EMBO Rep 5, 596–601 (2004).

71. Gordon, C.P. et al. Development of second-generation indole-based dynamin GTPase inhibitors. J Med Chem 56, 46–59 (2013).

72. Chen, Y., Gershlick, D.C., Park, S.Y. & Bonifacino, J.S. Segregation in the Golgi complex precedes export of endolysosomal proteins in distinct transport carriers. J Cell Biol 216, 4141–4151 (2017).

73. Sanjana, N.E. et al. A transcription activator-like effector toolbox for genome engineering. Nat Protoc 7, 171–192 (2012).

74. Cocucci, E., Gaudin, R. & Kirchhausen, T. Dynamin recruitment and membrane scission at the neck of a clathrin-coated pit. Mol Biol Cell 25, 3595–3609 (2014).

75. Ran, F.A. et al. Genome engineering using the CRISPR-Cas9 system. Nat Protoc 8, 2281–2308 (2013).

76. Zulkefli, K.L., Houghton, F.J., Gosavi, P. & Gleeson, P.A. A role for Rab11 in the homeostasis of the endosome-lysosomal pathway. Exp Cell Res 380, 55–68 (2019).

77. Curnock, R. & Cullen, P.J. Mammalian copper homeostasis requires retromer-dependent recycling of the high-affinity copper transporter 1. J Cell Sci 133 (2020).

78. McNally, K.E. et al. Retriever is a multiprotein complex for retromer-independent endosomal cargo recycling. Nat Cell Biol 19, 1214–1225 (2017).

79. Wang, P. et al. SNX17 Recruits USP9X to Antagonize MIB1-Mediated Ubiquitination and Degradation of PCM1 during Serum-Starvation-Induced Ciliogenesis. Cells 8 (2019).

80. Simonetti, B., Danson, C.M., Heesom, K.J. & Cullen, P.J. Sequence-dependent cargo recognition by SNX-BARs mediates retromer-independent transport of CI-MPR. J Cell Biol 216, 3695–3712 (2017).

81. Stockhammer, A. et al. Multi-functional ARF1 compartments serve as a hub for short-range cargo transfer to endosomes. bioRxiv, 2023.2010.2027.564143 (2023).

82. Edelstein, A., Amodaj, N., Hoover, K., Vale, R. & Stuurman, N. Computer control of microscopes using µManager. Curr Protoc Mol Biol Chapter 14, Unit14.20 (2010).

83. Schindelin, J., et al. Fiji: an open-source platform for biological-image analysis. Nat Methods 9, 676-682 (2012).

84. Ershov, D. et al. TrackMate 7: integrating state-of-the-art segmentation algorithms into tracking pipelines. Nat Methods 19, 829–832 (2022).

85. Tinevez, J.Y. et al. TrackMate: An open and extensible platform for single-particle tracking. Methods 115, 80–90 (2017).

86. Qiao, C. et al. Rationalized deep learning super-resolution microscopy for sustained live imaging of rapid subcellular processes. Nat Biotechnol 41, 367–377 (2023).

87. Shin, H.W., Morinaga, N., Noda, M. & Nakayama, K. BIG2, a guanine nucleotide exchange factor for ADP-ribosylation factors: its localization to recycling endosomes and implication in the endosome integrity. Mol Biol Cell 15, 5283–5294 (2004).

88. Temkin, P. et al. SNX27 mediates retromer tubule entry and endosome-to-plasma membrane trafficking of signalling receptors. Nat Cell Biol 13, 715–721 (2011).

